# Insights into Membrane Protein-Lipid Interactions from Free Energy Calculations

**DOI:** 10.1101/671750

**Authors:** Robin A. Corey, Owen N. Vickery, Mark S. P. Sansom, Phillip J. Stansfeld

## Abstract

Integral membrane proteins are regulated by specific interactions with lipids from the surrounding bilayer. The structures of protein-lipid complexes can be determined through a combination of experimental and computational approaches, but the energetic basis of these interactions is difficult to resolve. Molecular dynamic simulations provide the primary computational technique to estimate the free energies of these interactions. We demonstrate that the energetics of protein-lipid interactions may be reliably and reproducibly calculated using three simulation-based approaches: potential of mean force calculations. alchemical free energy perturbation, and well-tempered metadynamics. We employ these techniques within the framework of a coarse-grained force field, and apply them to both bacterial and mammalian membrane protein-lipid systems. We demonstrate good agreement between the different techniques, providing a robust framework for their automated implementation within a pipeline for annotation of newly determined membrane protein structures.

## Introduction

Integral transmembrane proteins have diverse functions within cells, and as such are key targets for many drugs, ranging from antibiotics to anticancer agents. Structurally, they are unified by the presence of a hydrophobic span of residues that both anchors the protein within the core of a lipid bilayer membrane and presents the flanking residues to the surrounding polar lipid head groups. The resulting protein-lipid interactions are important for function, with many membrane proteins, including e.g. ion channels, transporters and receptors, regulated by specific lipid interactions ^1^. Lipid-binding sites thus provide potential druggable allosteric sites on many biologically-important membrane proteins.

Structural studies of membrane proteins often rely on their extraction from their native bilayer environment through use of detergents. As a consequence of this, lipids which bind to the protein are often lost before structural (X-ray diffraction or cryoelectron microscopy) data are gathered. Although there are cases where X-ray or electron scattering density may be observed for lipids bound to membrane proteins (for examples, see refs ^2–4^), the often modest resolution of such data presents challenges to the unambiguous assignment of the molecular identity of the bound lipid species.

Molecular simulations provide near atomic resolution insights into the interactions of lipids with membrane proteins. They can both predict the location of lipid binding sites in advance of structural studies ^5–7^ and can extend structural observations on the lipid interactions of a given membrane protein to other members of a protein family ^8^. In addition to identification of potential lipid interaction sites, for example from estimates of lipid-protein ‘fingerprints’ ^9^, molecular simulations can provide estimates of the residence times of lipids at binding sites on a membrane protein ^10^ and of free energies of interaction of specific lipids ^11,12^.

Validation of computational predictions of specific lipid interactions can be achieved via a number of biophysical approaches, including e.g. native mass spectrometry (nMS) ^13^ which can be employed in tandem with molecular simulation ^14^. The relatively slow throughput of these techniques however, means that only a tiny fraction of the possible interactions has so far been identified. Moreover, experimental quantification of the strength and specificity of protein-lipid interactions remains more challenging, with notable recent attempts using nMS ^15^ and surface plasmon resonance (SPR)-based methods ^16^.

Molecular simulations can also be used to quantify the strength of protein-lipid interactions, via free energy calculations (Figure 1A). Several free energy techniques have been developed for the calculation of binding free energies between ligands and (water soluble) proteins ^17^, including online tools for ease of set-up, e.g. *pmx* (http://pmx.mpibpc.mpg.de/webserver.html). Many, if not all, of these methods can be modified for analysis of protein-lipid interactions. Membrane proteins and lipids pose particular challenges of sampling and convergence for accurate free energy estimation ^18^, arising from the relatively slow rates of lipid diffusion and from the diversity of lipid species present in complex biological membranes ^19^. To date, most studies ^5,11,18,20,21^ have combined umbrella sampling with a potential of mean force (PMF) calculation along a one-dimensional reaction coordinate connecting the binding site with the surrounding membrane ^18^ (Figure 1B). Convergence of such calculations (i.e. the point at which additional sampling via additional simulation does not substantially change the outcome) is often achieved through use of a coarse-grained (CG) biomolecular force field, such as Martini ^22,23^, which allows for efficient sampling of protein-lipid interactions. Whilst a powerful technique, the difficulty in demonstrating convergence makes this approach challenging to implement in a high throughout fashion. Furthermore, it is computationally demanding, requiring ~50+ μs of simulation per protein-lipid interaction, currently equivalent to ~2 weeks on a typical GPU-node. It is therefore important that we explore additional approaches in order to extend the reach and to evaluate the robustness of PMF-based estimates of free energies of protein-lipid interactions.

**Figure 1.**
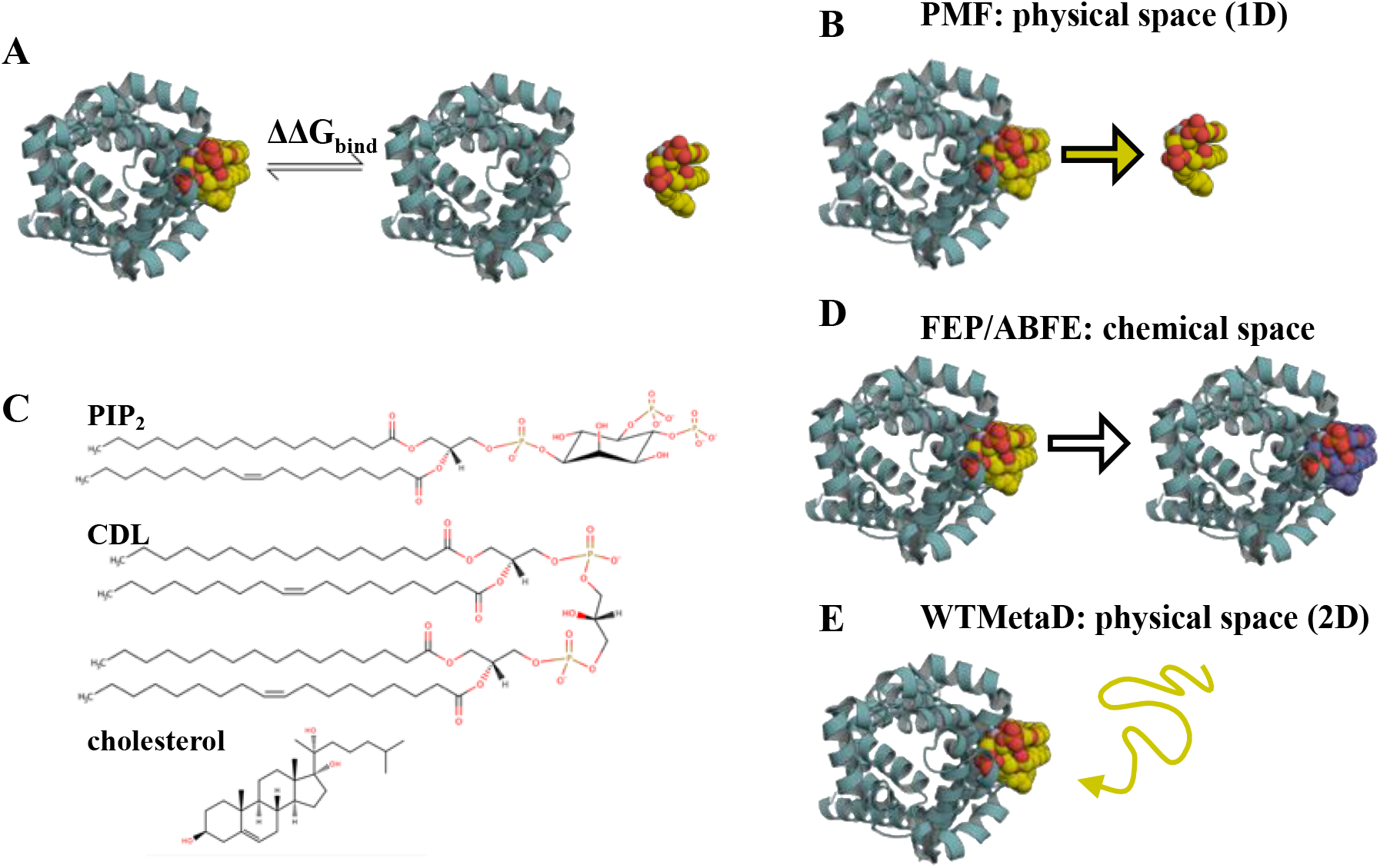
Introduction to free energy calculations. A) Overview of free energy calculations. A membrane protein, as viewed from above the membrane, is shown in cyan cartoon, and a lipid in yellow, orange and red spheres. Two states are modelled: the left state is the protein bound to the target lipid, the right is the protein bound to a generic lipid (not shown), with the target lipid unbound. Free energy calculations aim to compute the difference in free energy between these states (ΔΔG_bind_). B) Potential of mean force (PMF) calculations create a reaction coordinate in physical space by pulling the lipid away from or towards the binding site. This coordinate can then be sampled, e.g. with umbrella sampling, to provide a 1D energetic landscape, allowing calculation of ΔΔG_bind_ between the target and a generic lipid. C) Chemical structures of PIP_2_, cardiolipin (CDL) and cholesterol. D) Free energy perturbation (FEP) and absolute binding free energy (ABFE) calculations build alchemical pathways which either change the bound lipid into a different species, in this study to that of the bulk membrane, or fully remove the lipid from the simulation box. This provides the binding free energy difference between the target lipid and a generic lipid, ΔΔG_bind_. E) Well-tempered metadynamics (WTMetaD) biases the diffusion of a target lipid around the protein through addition of a time-dependent Gaussian of energy to the collective variable (CV). These Gaussians can then be reconstructed into a full 2D energy landscape, with comparison of binding regions and the bulk membrane giving ΔΔG_bind_.

Here, we present an analysis of the determination of protein-lipid binding interactions using PMFs alongside two other powerful free energy approaches, adapted here for investigation of protein-lipid interactions. These are free energy perturbation (FEP ^24^) or absolute binding free energy (ABFE ^25^) calculations, whereby a molecule is partially or fully perturbed via non-natural (i.e. alchemical) chemical space (Figure 1D), and well-tempered metadynamics (WTMetaD ^26^), where a history-dependent bias is added to a free energy surface (Figure 1E) to reduce simulation time spent sampling local energy minima. We compare PMF, FEP/ABFE and WTMetaD in terms of ease of accuracy and computational cost. We use all three methods on a panel of experimentally well-characterised proteins which are representative of bacterial, mitochondrial and mammalian cell membranes. Through comparison of the methods applied here we outline a mechanism for producing robust and reproducible estimates of protein-lipid interactions from molecular simulations.

## Methods

### Equilibrium CG simulations

Simulations were run using the CG Martini v2 biomolecular force field ^22,23^. In this forcefield, molecules are coarse grained through the representation of approximately 4 heavy atoms and associated hydrogens as a single bead or particle. Whilst this simplification provides the force field with reduced resolution ^27^, it has repeatedly been shown as highly proficient in the identification of specific interactions between proteins and membrane lipids, including cardiolipin (CDL) ^11,21,28,29^, phosphatidylinositol (4,5) bisphosphate (PIP_2_) ^5,14,20^, cholesterol ^20,30^ (Figure 1C) and others.

We follow the MemProtMD protocol for setting up CG simulations of integral membrane proteins in an equilibrated bilayer ^31,32^. Briefly, the input proteins are aligned accordingly on the *xy* plane, and lipids (1-palmitoyl-2-oleoyl-sn-glycero-3-phosphocholine; POPC or 1-palmitoyl-2-oleoyl-sn-glycero-3-phosphoethanolamine; POPE) are placed randomly around the transmembrane region of protein, in a *z* range of 8 nm. The starting protein coordinates were used as follows: chicken (*Gallus gallus*) Kir2.2 ion channel, PDB code 3SPC ^33^; bovine (*Bos taurus*) AAC transport protein, PDB code 1OKC ^4^; bacterial (*Aquifex aeolicus*) LeuT transport protein, PDB code 2A65 ^34^ (simulated here as a dimer); and human (*Homo sapiens*) GPCR A_2A_R, PDB code 5IU4 ^35^ with the bRIL subunit removed. In each case, non-protein atoms were removed and any missing loops were added using MODELLER ^36^ or SWISS-MODEL ^37^ prior to CG conversion. Phospholipids were modelled with palmitoyl (4-beads) and oleoyl (5-beads) tails. Cholesterol was modelled using the virtual-site description ^38^. Full details of the simulations are given in the Supporting Methods.

### Potential of mean force calculations

PMF calculations were set up and run as described previously ^18^. Calculations start from the complex formed between the protein and target lipid, which is then inserted into a generic membrane (i.e. POPE or POPC). The lipid is then removed from the protein through application of steered MD, in which a distance-dependent pulling force is applied between the lipid head group and the protein. This trajectory then forms the collective variable (CV) to be analysed. The specific CVs used here are outlined in the Supporting Methods.

Snapshots were taken of the system in which the lipid is at specific window along the CV (using a 0.05 nm spacing for optimal histogram overlap, see e.g. Supporting Figure 1), with each snapshot used to seed an independent simulation. For these, an umbrella potential with a force of 1000 kJ mol^-1^ was used to keep the lipid in place along the reaction coordinate, with 100 kJ mol^-1^ nm^-2^ *xy* positional restraints on 3-4 protein backbone beads to prevent the protein from rotating during the simulations – details provided in the Supporting Methods.

Simulations were run for 1 μs to allow convergence (e.g. Supporting Figure 1). The first 200 ns were removed from each simulation as equilibration, and the final 800 ns were combined into a 1D energy profile using the weighted histogram analysis method (WHAM) ^39^, as implemented in *gmx wham* ^40^, and employing 200 rounds of Bayesian bootstrapping to report on statistical accuracy. When plotting, the bulk region of the membrane is considered to have a free energy of 0 kJ mol^-1^, and the binding energy well is set to 0 nm on the *x*-axis.

Note that, as the lipid binding site will either be occupied by the target lipid or a generic bulk lipid, this analysis will not provide us with ΔG_bind_ of our target lipid to the site, but instead ΔΔG_bind_ between the target and a generic lipid. This is the biologically appropriate term, as protein-lipid binding will always occur in competition with other lipids, and the effective affinity of the interaction will be dependent on the nature of the other lipids present. Accordingly, if you carry out PMF calculations of e.g. POPC in a POPC membrane, the value reported should be 0 kJ mol^-1^ (e.g. see ref ^21^).

### Free energy perturbation

The bound PMF systems were used as the input for the FEP calculations. The target lipid was alchemically transformed into the generic lipid, along a coordinate in chemical space, termed λ. Additional simulations were run perturbing the target lipid in bulk membrane (‘free’), i.e. with no protein visible to the lipid. ΔΔG_bind_ can then be calculated as described in Figure 2. This value should be equivalent to that obtained using PMF calculations.

**Figure 2.**
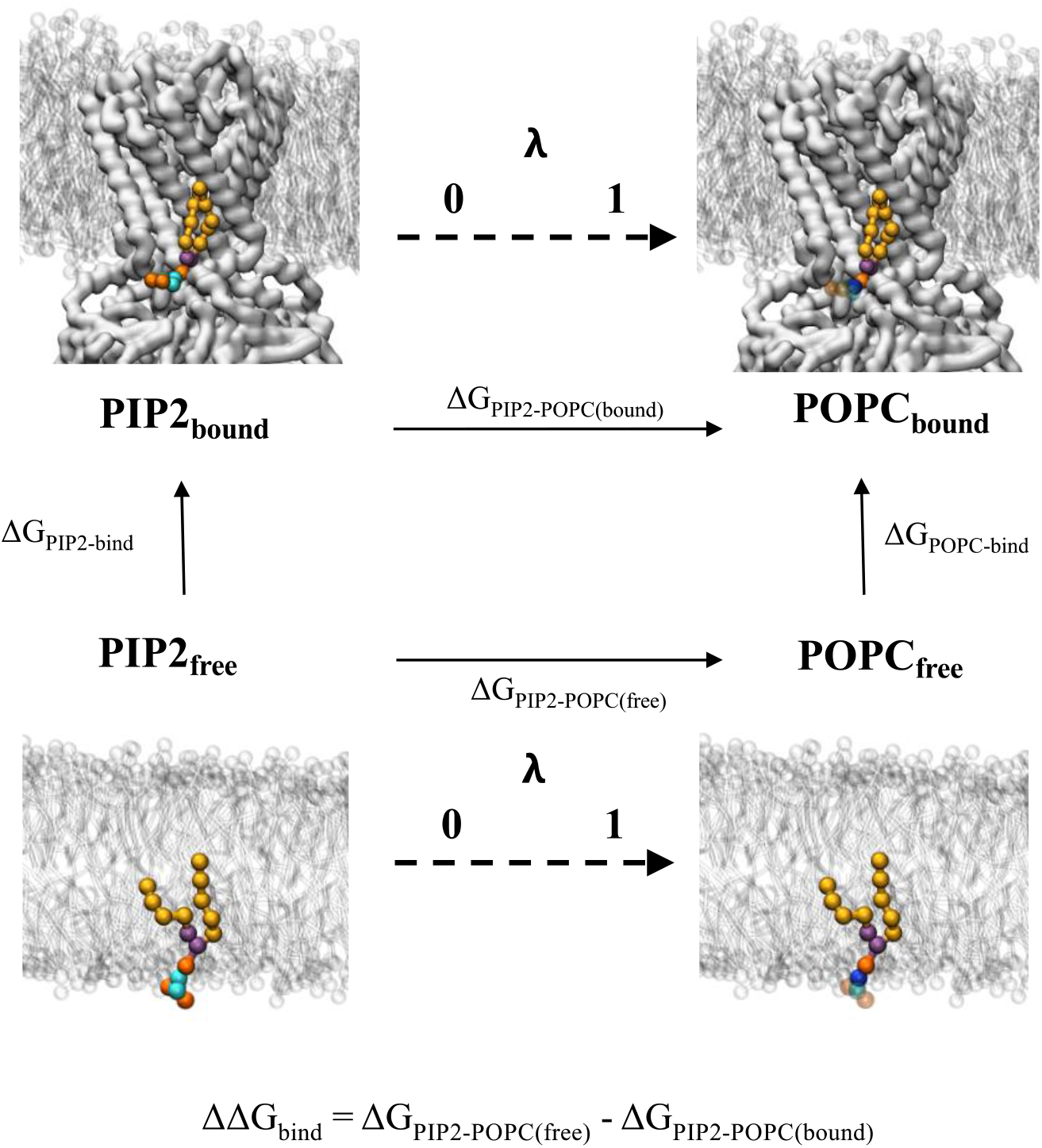
Thermodynamic cycle for FEP. Shown is a representative thermodynamic cycle used for the FEP calculations in this study. The protein receptor (here Kir2.2) is shown as white surface, with the membrane as white sticks. The target lipid is shown as coloured spheres: on the left the native PIP_2_ molecule has lipid tails in yellow, glycerol beads in purple, phosphate beads in orange and sugar beads in cyan. On the right, the perturbed POPC lipid is coloured as PIP_2_, but with a blue choline bead, and transparent beads for the beads which have now been decoupled from the system. The horizontal vertices represent the two FEP calculations, where the target lipid is alchemically perturbed into the bulk lipid either free in membrane (ΔG_PIP2-POPC(free)_) or bound to the receptor (ΔG_PIP2-POPC(bound)_). These reactions are represented by a λ coordinate from 0 to 1. ΔΔG_bind_ can be calculated as ΔG_PIP2-POPC(free)_ - ΔG_PIP2-POPC(bound)_, which will represent the same value as calculated in the PMF calculations (Figure 1). Note that the right vertical vertex (ΔG_POPC-bind_) is the free energy required for the bulk lipid binding the receptor – in this special case of protein-lipid binding, where the ligand and the solvent are the same molecule, this value should be 0 kJ mol^-1^.

Charges and Lennard Jones interactions were turned off separately, with a soft-core parameter (sigma=0.3) used for the Lennard Jones interactions ^41,42^. Bonded interactions were not perturbed, as these cancel out between the bound and free states. For each system, we used a single topology method, where molecule A is converted into molecule B in a single transformation. Details of the transformation are provided in the Supporting Methods. In all cases, we used 10 windows to perturb the Columbic interactions, and either 10 (PIP_2_) or 20 (CDL) windows were used to perturb the Lennard-Jones interactions.

Each λ window was minimized using the steepest descent method, followed by 5 ns of NPT equilibrium. Five independent production simulations of 250 ns were then run with randomized initial velocities, using a leap-frog stochastic dynamics integrator.

The free energy pathway was then constructed from the individual λ windows using the Alchemical Analysis package ^43^. Energy values were calculated on the final 225 ns of simulation data, with this simulation length showing good conversion (e.g. Supporting Figure 2). Analysis was run using the MBAR method ^44^, although we observed good agreement between multiple analysis methods (e.g. Supporting Figure 2). Data from 5 repeats were averaged, and the standard deviations calculated.

### Absolute binding free energy calculations

A_2A_R-cholesterol simulations followed a similar setup as described above, but with the target lipid fully, rather than partially, decoupled from the simulation box (Supporting Figure 3). To keep the cholesterol molecule in the binding site and within the correct plane and orientation of the membrane at high values of λ, we followed a cholesterol restraining scheme described previously ^45^, and outlined in the Supporting Methods. ΔΔG_bind_ was calculated from the energy required to decouple the cholesterol molecule in the bound and free state, in each case perturbing the Lennard-Jones interactions over 29 λ windows (with 0.05 spacing from 0 to 0.6, and 0.025 spacing from 0.6 to 1). The energetic input of the restraints accounted for through two analytical terms (see Supporting Equations 1, 2 and 3), and additional restraint FEP in the gas phase (see Supporting Figure 3). Simulations were run in Gromacs 2016 (www.gromacs.org) with restraints imposed using the *plumed* v2.4 plugin ^46,47^.

### Metadynamics

In classical MetaD, the evolution of the system along the CV is biased by a history dependant potential, which is the sum of the Gaussians deposited along the relevant CV ^48^. After a defined period of time, the biasing potential compensates the underlying Free Energy Surface (FES), allowing the real FES to be estimated. Use of a fixed Gaussian height gives classical MetaD several limitations, particularly concerning convergence, as the system can be pushed into regions of configurational space which are not physically or physiologically relevant. Here we use Well-Tempered MetaDynamics (WTMetaD) ^26,49^ in which the Gaussian height is rescaled based upon an adaptive bias (ΔT), which is dependent on the system simulated ^26^.

The WTMetaD simulations were made using the protein coordinates built into a simple (POPC or POPE) membrane as described above, with *xyz* positional restraints on select backbone particles – details provided in the Supporting Methods. The WTMetaD simulations were run using 20 walkers placed randomly within 2.5 nm of the protein. In the A_2A_R, AAC and Kir2.2 systems, the walkers were constrained from moving greater than 3.5, 4 and 4.2 nm respectively from the geometric centre of the protein. For LeuT the walkers were constrained using a minimum distance of 4 nm between the lipid and the geometric centre of each monomer. To prevent the cholesterol from flip-flopping between leaflets, two additional flat-bottomed restraints were applied: an angle restraint of one radian was applied between the ROH, C2 beads and the *z*-axis and a wall parallel to the midpoint of the membrane offset towards the extracellular leaflet by 1 nm was applied to the ROH bead to prevent the cholesterol from moving vertically between leaflets. In all simulations the flat bottom restraints were applied using the Plumed UPPERWALLS of 1,000 kJ mol^-1^ nm^-2^.

The bias was added along the CV for each system, as defined in the Supporting Methods. The bias-factor was tuned to each lipid with cholesterol, CDL and PIP_2_ being 6, 8, and 20 respectively. The following WTMetaD parameters were applied to all systems: Gaussian width of 0.01 nm, height of 1 kJ mol^-1^ and a 1 ps deposition rate. All WTMetaD simulations were performed at 310 K using Gromacs 2016 with the Plumed v2.4 plugin. When plotting, the bulk region of the membrane is considered to have an energy of 0 kJ mol^-1^.

The bulk Gaussian height of 5 % of the maximum value was used as a metric for the WTMetaD reaching a steady state (e.g. Supporting Figure 4), at which we see multiple association/disassociation events to the binding sites. The FES was sampled every 250 ps, whereupon the final 2D FES depicted in the text was recovered by averaging over the steady state.

## Results

We selected a panel of four experimentally well-characterised membrane proteins to represent bacterial, mitochondrial, and mammalian cell membranes. These were chosen to include examples for which experimental structural data for protein-lipid interactions are available, for which nMS studies have been used to demonstrate lipid interactions, and which have been the subject of previous computational studies. They also represent three species of lipid (Figure 1C) which frequently form interactions with membrane proteins, namely: PIP_2_, a negatively charged phospholipid which interacts with many ion channels and receptors in mammalian cell membranes; CDL, also negatively charged, which is present in mitochondrial and bacterial inner membranes, and cholesterol, which has been observed to bind to many G-protein coupled receptors (GPCRs) and ion channels. Thus, we probe the following interactions: of PIP_2_ with the mammalian inward rectifying potassium channel, Kir2.2; of CDL with the mitochondrial inner membrane ADP/ATP carrier protein, AAC ^21^ and with the bacterial inner membrane leucine transporter, LeuT ^34,50^; and of cholesterol with a GPCR, the adenosine 2A receptor (A_2A_R).

### Kir2.2-PIP_2_ binding interactions

Kir2.2 is a member of the inwardly rectifying potassium channel family, found in neuronal cell membranes, which play a key role in the regulation of plasticity and neuronal excitation ^51^. As with other members of this family, opening of Kir2.2 channels can be activated through interaction with PIP_2_ ^52^. Structural (X-ray) studies show PIP_2_ binds at the interface between the transmembrane and cytoplasmic domains (Figure 3A) to bring about channel opening ^33^.

**Figure 3.**
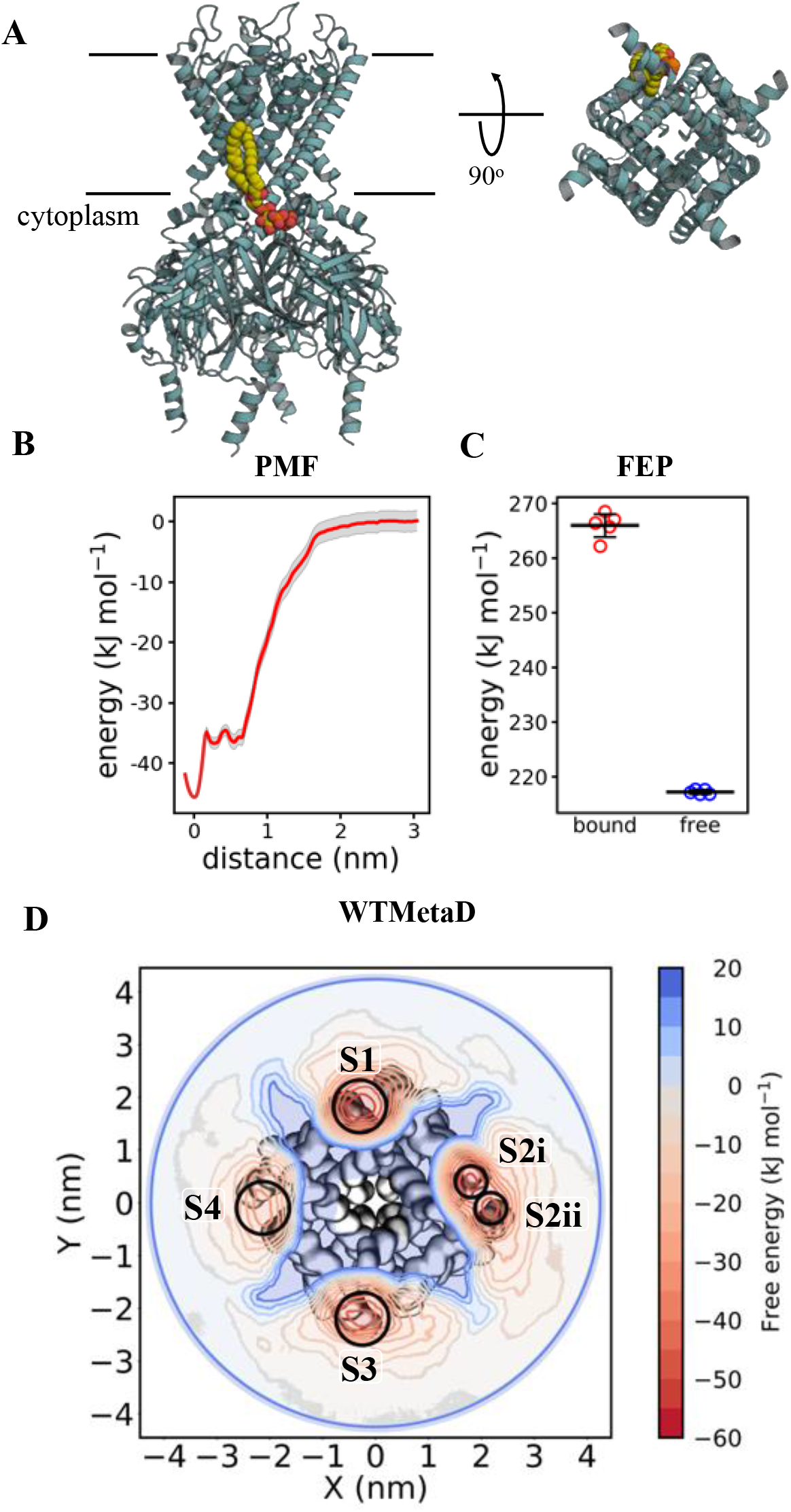
Calculating Kir2.2-PIP_2_ binding energetics. A) View of Kir2.2 in cyan cartoon, with a bound PIP_2_ molecule in yellow, orange and red spheres, as sampled with CG MD. PIP_2_ interactions map to residues Arg 78, Arg 80, Lys 183, Arg 186, Lys 188, Lys 189 (*Gallus gallus* numbering). The approximate position of the membrane is shown with black lines. On the left is a view from the side, and on the right is a cytoplasmic view of the transmembrane region alone, with the intracellular domain removed for clarity. Note that only one PIP_2_ binding pose is shown, but four are present around the homotetrameric Kir2.2. B) PMF data for Kir2.2-PIP_2_ binding. The *y*-axis is set to 0 for the bulk membrane, and the difference between this and the energy well (set to 0 nm on the *x*-axis) is ΔΔG_bind_, here −45±2, with errors from 200 rounds of bootstrap analysis. C) FEP data for Kir2.2-PIP_2_ binding, showing the energy cost for perturbing PIP_2_ to POPC whilst bound to Kir2.2 (red) and whilst free in a POPC membrane (blue). ΔΔG_bind_ can be calculated from the free data minus the bound (see Figure 2), giving a value of −48±2, with the error the standard deviation from five repeats. D) 2D energy landscape for Kir2.2 and PIP_2_ as computed using WTMetaD. The protein is shown as surface behind the data, with the large intracellular domain removed for clarity. The energetic landscape for a PIP_2_ molecule around the protein has been computed and is shown as a red-blue contour map. Four binding regions in red can be seen around the protein, with reported ΔΔG_bind_ values as follow: S1=-55±7, S2i=-49±4, S2ii=-45±5, S3=-45±6 and S4=-36±7 kJ mol^-1^.

Simulations were run to probe the free energy of Kir2.2-PIP_2_ interactions in a simple phospholipid (POPC) membrane. Binding sites have previously been identified both structurally ^33^ and computationally ^5,18^. We, therefore, carried out PMF and FEP simulations based on the crystallographic binding pose of PIP_2_ in order to estimate the binding free energy. We note that these PMFs provide an interaction free energy for Kir2.2 with PIP_2_ relative to that with POPC, i.e. the well depth in the PMF can be equated as ΔG_bind-PIP2_ - ΔG_bind-POPC_, or ΔΔG_bind(PIP2-POPC)_, which we shorten to ΔΔG_bind_. Similarly, the FEP analysis provides us with ΔΔG_bind_ (see Figure 2).

The PMF and FEP data agree well, giving similar estimates for Kir2.2-PIP_2_ interactions (Figure 3B-C; - 45±2 kJ mol^-1^ for PMF and −48±2 kJ mol^-1^ for FEP). The PMF data reveal the presence of a second binding site, ~0.5 nm from the main site, with a free energy of interaction of −36±2 kJ mol^-1^. This alternative pose of the lipid involves interactions between PIP_2_ and residue Lys 220, and corresponds to a secondary binding mode proposed for the closely related Kir2.1 channel ^53^.

Next, we carried out WTMetaD simulations to explore the multiple PIP_2_ binding sites on the tetrameric channel structure. As before, the WTMetaD data allows us to calculate ΔΔG_bind_ of PIP_2_ in relation to POPC. The data identified the primary binding modes for three of the channel subunits, with interaction free energies of −55±7 kJ mol^-1^, −49±4 kJ mol^-1^ and −45±6 kJ mol^-1^ (Figure 3D). The latter of these values corresponds to the site probed using PMF and FEP, revealing excellent agreement between the techniques. For the fourth subunit, the secondary binding mode was recovered (−36±7 kJ mol^-1^), which again corresponds well with value from the PMF analysis.

We note that in the WTMetaD simulations of Kir2.2, where the protein is necessarily *xyz* restrained (see Supporting Methods), the channel is tilted (by ca. 5°) relative to the bilayer normal. We expect that this small tilt relative to the bilayer induces the shift of PIP_2_ from the primary to the secondary binding site for the fourth channel subunit, with the primary binding mode only accessible to three of the four binding sites due to the orientation of the channel.

### CDL interactions with two transport proteins

The ADP/ATP carrier (AAC), also known as the adenine nucleotide translocator (ANT), is present in the inner mitochondrial membrane where it accounts for 10% of the total protein content ^54^. It functions as a regulator of mitochondrial adenine nucleotide concentration, allowing flux of ATP/ADP across the mitochondrial membrane ^55^. CDL is known to bind to AAC ^4,56^ (Figure 4A), where it results in activation of the transporter ^57,58^. The protein has an approximate three-fold symmetry with three homologous, but potentially non-identical, binding sites for CDL.

**Figure 4.**
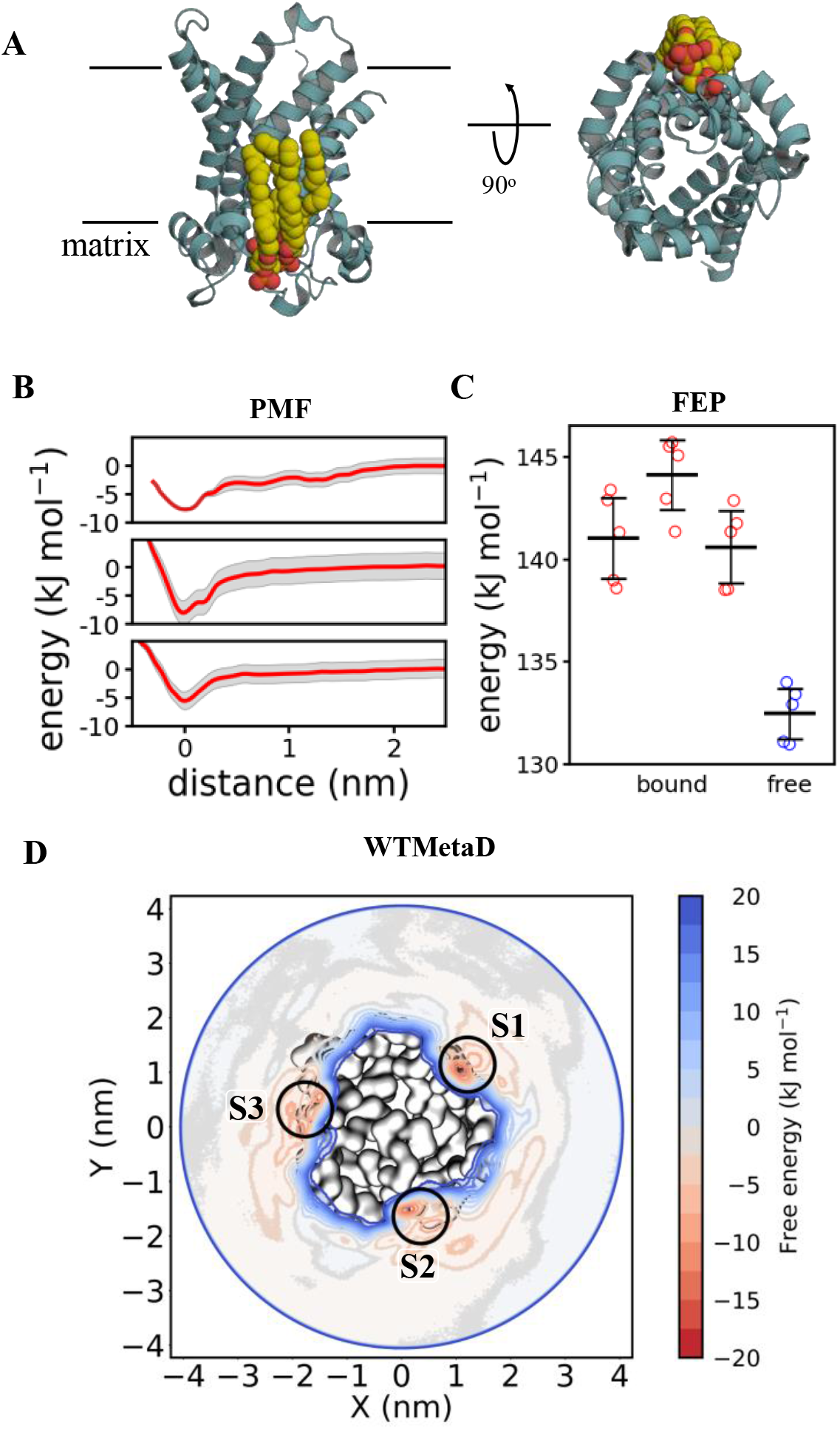
Calculating AAC-CDL binding energetics. A) AAC bound to CDL. Colours and views as in Figure 3A. Note that three CDL binding sites are present around AAC, which is a homotrimer. B) PMF data for each of the AAC-CDL binding sites, as per Figure 3B. Site 1 is at the top, site 2 in the middle and site 3 at the bottom. ΔΔG_bind_ is −7±2, −8±4 and −5±2 for each site respectively. C) FEP data for AAC-CDL binding, showing the energy cost for perturbing CDL to POPC whilst bound to AAC in each of the three binding sites (red: site 1 on the left) and whilst free in a POPC membrane (blue). ΔΔG_bind_ is −9±3, −11±3 and −8±2 for each site respectively. D) 2D energy landscape for AAC and CDL, as per Figure 3D. Three binding regions in red can be seen around the protein, reporting ΔΔG_bind_ values as follow: S1=-14±3, S2=-13±3 and S3=-11±6 kJ mol^-1^.

We probed the energetics of CDL-AAC interaction at all three of these sites with the protein embedded in a POPC membrane using PMF and FEP calculations (Figure 4A). PMF analyses yielded free energies of −7±2, −8±4 and −5±2 kJ mol^-1^ for CDL binding at these sites (Figure 4B). FEP produces good agreement with the PMF data, giving −9±3, −11±3 and −8±2 kJ mol^-1^ (Figure 4C), with each technique ranking the sites the same in terms of CDL binding energies.

Next, we explored the system using WTMetaD, which revealed clear energy wells at all three binding sites (Figure 4D). The interaction free energies for sites 1, 2 and 3 were −14±3 kJ mol^-1^, −13±3 kJ mol^-1^, and −11±6 kJ mol^-1^ respectively, showing reasonable agreement with the PMF and, to a greater degree, FEP estimates. Reassuringly, the sites are again similarly ranked, with the site 3 lower in energy than sites 1 or 2.

We also explored CDL interactions with the leucine transporter (LeuT), which is a bacterial homologue of the solute carrier family 6 (SLC6) class of proteins, which includes the human serotonin transporter ^59^. LeuT catalyses sodium-driven small hydrophobic amino acid transport across the bacterial inner membrane ^60^, and has been shown to bind to CDL (Figure 5A), which stabilizes the dimeric form of the transporter ^50^. We probed the energetics of the CDL-LeuT interaction in a POPE membrane using both PMF and FEP calculations. The initial CDL pose was based on a previously-identified likely binding site at the dimer interface (see ref ^50^ for more details). PMF analysis yielded a value of −6±3 kJ mol^-1^ for CDL at this site (Figure 5B), in agreement with the FEP value of −9±2 kJ mol^-1^ (Figure 5C). WTMetaD not only identifies the dimer interface binding site (with a binding free energy of −7±3 kJ mol^-1^ and −10±6 kJ mol^-1^ for the equivalent and opposing sites respectively), but also additional sites around the complex (Figure 5D). For example, sites 2 and 4 appear to bind CDL with a lower energy than the dimer-interface site (−3±2 kJ mol^-1^ for each site). Non-specific of CDL with a low free energy of ca. −3 kJ mol^-1^ is also seen over a 3 nm area of each monomers (Figure 5D; dotted lines).

**Figure 5.**
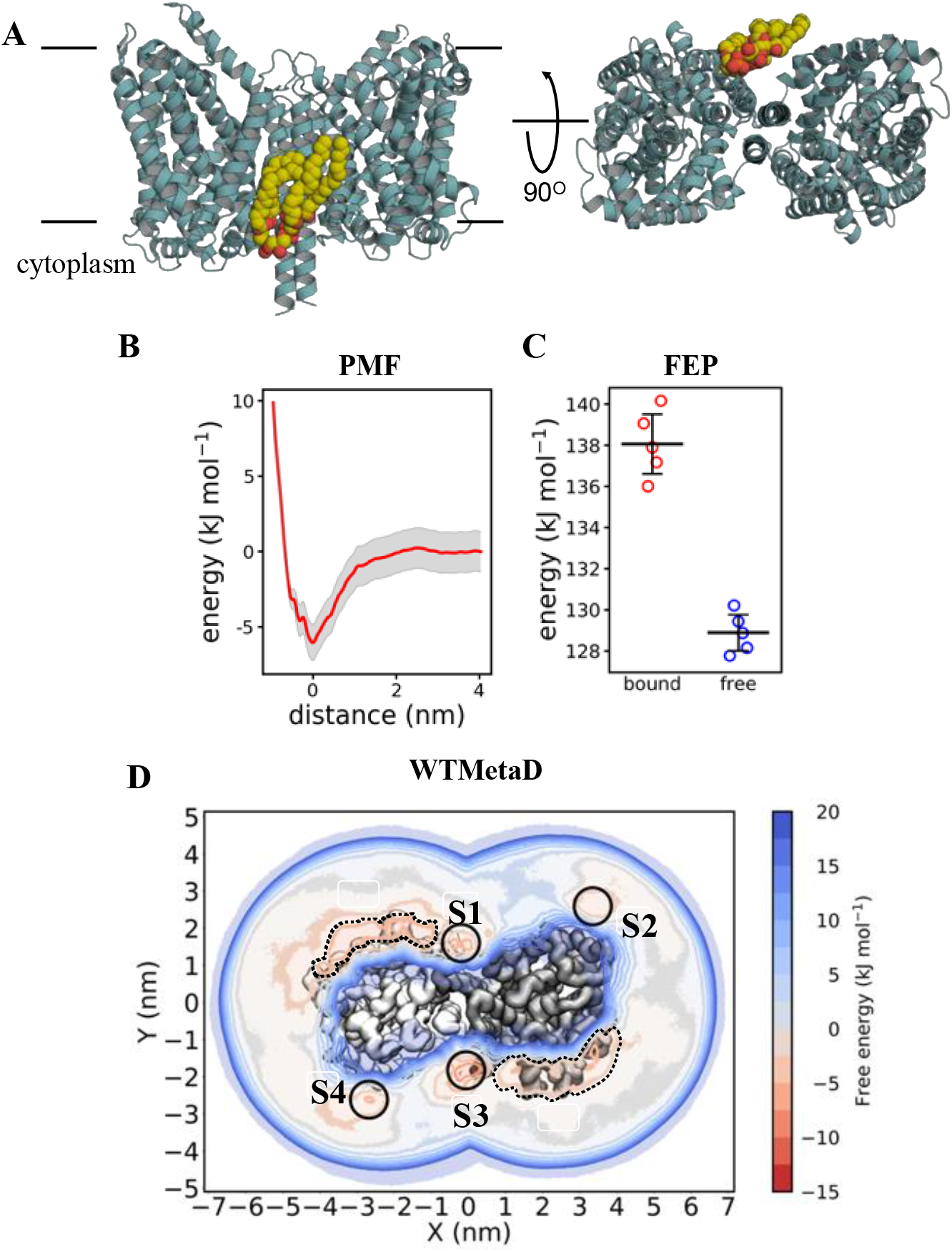
Calculating LeuT-CDL binding energetics. A) View of LeuT bound to CDL. Colours and views as in Figure 3A. Note that a second equivalent CDL binding site is present on the other side of the homodimeric LeuT. B) PMF data for AAC-LeuT binding, as per Figure 3B. ΔΔG_bind_ is −6±3. C) FEP data for AAC-LeuT binding, perturbing CDL to POPE whilst bound to LeuT (red) and whilst free in a POPE membrane (blue). ΔΔG_bind_ is −9±2. D) 2D energy landscape for LeuT and CDL, as per Figure 3D. Two binding regions in red can be seen around the protein, reporting ΔΔG_bind_ values as follow: S1=-7±3, S2=-3±2, S3=-10±6 and S4=-3±2 kJ mol^-1^ Two indiscriminate binding regions of ca. −3 kJ mol^-1^ are highlighted with dotted lines.

### Cholesterol interactions with a GPCR

The G-protein coupled receptor (GPCR) class of proteins constitute the largest family of membrane proteins and account for 35% of all drug targets ^61^. Many members are thought to be functionally modulated by cholesterol binding ^62^, including the highly studied adenosine 2A receptor (A_2A_R). This provides an example of protein-lipid interactions with an uncharged and relatively rigid lipid molecule.

Cholesterol binding to A_2A_R has been previously identified structurally ^63^ and explored using molecular dynamics ^7,20,64^ To probe the energetics of this process, we applied WTMetaD to human A_2A_R in a POPC membrane to produce a full 2D map of cholesterol binding on the extracellular leaflet around the receptor. We identified three binding sites between TM 1-2, 1-7 and 6-7 (Figure 6A-B), with binding free energies of −8±3, −9±3 and −5±2 kJ mol^-1^. Of these only the TM 6-7 cholesterol binding site has been captured in structural data for A_2A_R ^35,65,66^, although the other two binding sites correspond to electron density identified as acyl tails in several A2AR structures ^35,65,66^.

**Figure 6.**
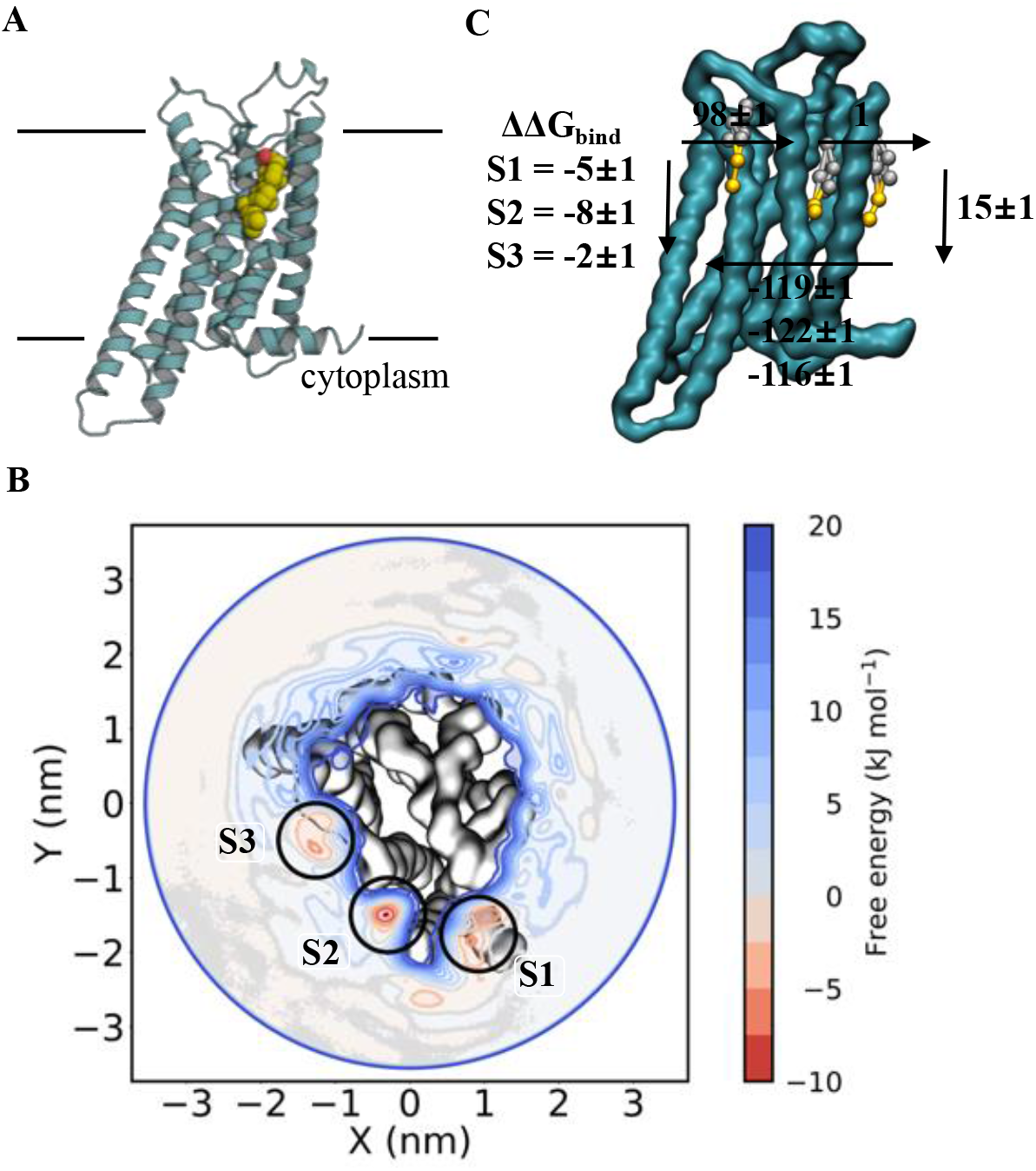
Calculating A2AR-cholesterol binding energetics. A) View of A2aR bound to cholesterol, in one of several possible binding sites. Colours as in Figure 3A. B) WTMetaD 2D energy landscape for A_2A_R and cholesterol, for the extracellular leaflet only. Three binding regions in red can be seen around the protein, reporting ΔΔG_bind_ values as follow: S1=-8±3, S2=-9±3 and S3=-5±2 kJ mol^-1^. C) ABFE analysis the sites from panel B. Here, the bound cholesterol is fully decoupled from a POPC membrane whilst bound to A_2A_R or whilst free in the membrane. The cycle represented here is a simplified version of the cycle in Supporting Figure 3. The final ΔΔG_bind_ values are −5±1, −8±1 and −2±1 kJ mol^-1^.

We then probed these site using ABFE, using restraints and a thermodynamic cycle outlined in a recent study by Salari et al ^45^ (see Supporting Methods and Supporting Figure 3). We obtained values of −5±2 kJ mol^-1^, −8±1 kJ mol^-1^ and −2±1 kJ mol^-1^ for the three sites (Figure 6C), in reasonable agreement with the WTMetaD data. Importantly, we get an identical ranking of the sites between the techniques, with site 2 the highest energy and site 3 the lowest. Note that we were unable to probe these sites using PMF calculations, as the energies become swamped by background thermal fluctuations.

We can compare our estimates of the strength of A2AR cholesterol interactions with those from other simulation studies of GPCRs. Genheden et al ^67^ estimated free energies of interaction with b2AR and A_2A_R of the order of −10 to −15 kJ mol^-1^ from CG simulations. Lee and Lyman ^7^ estimated free energies for the A2aR of −3 to −5 kJ mol^-1^ from atomistic MD simulations. Thus, our estimates are in broad agreement with those from previous studies of the A_2A_R, both of which estimated free energies directly from cholesterol occupancies following extended (but possibly under sampled) equilibrium MD simulations.

## Discussion

Integral membrane proteins are strongly affected by the lipid environment in which they reside (for a recent review, see ref ^1^). Information on the structural basis of these interactions can be determined by X-ray crystallography, cryoelectron microscopy, and NMR. In contrast, determining the free energies of these interactions remains very challenging either experimentally ^15,16,29^, or computationally ^5,11,18,20,21^.

One of the main challenges facing computational estimation of free energies of protein-lipid interactions is presented by the slow timescale of relaxation of lipid molecules in a bilayer. This means that for free energy calculations, we need to be confident that we have adequately sampled those interactions, i.e. that the simulations have converged. The agreement between all three techniques for lipid binding energetics gives us confidence that this is indeed the case (Table 1 and Figure 7). To our surprise, the analyses here provide lower energies for CDL binding to AAC ^21^ and other proteins ^11,18,68^ than previously reported; this discrepancy most likely reflects a different handling of long range electrostatic interactions. Nonetheless, when comparing like-for-like systems, as here, it is evident that all three techniques converge to a common value and provide a similar degree of accuracy.

**Table 1.** Summary of binding energies for the systems tested here. Reported are the measured ΔΔG_bind_ values, with the errors calculated as defined in the text and figure legends. These data have been used to populate the chart in Figure 7.

| system | site | lipid | membrane | PMF | FEP/ABFE | WTMetaD |
| --- | --- | --- | --- | --- | --- | --- |
| Kir2.2 | 3 | PIP<sub>2</sub> | POPC | -45 ± 2 | -48 ± 2 | -45 ± 6 |
| AAC | 1 | CDL | POPC | -7 ± 2 | -10 ± 4 | -14 ± 3 |
|  | 2 |  |  | -8 ± 4 | -11 ± 2 | -13 ± 3 |
|  | 3 |  |  | -7 ± 4 | -9 ± 4 | -11 ± 6 |
| LeuT | 1 | CDL | POPE | -6 ± 3 | -9 ± 2 | -7 ± 3 |
| A2AR | 1 | chol | POPC |  | -5 ± 1 | -8 ± 3 |
|  | 2 |  |  |  | -8 ± 1 | -9 ± 3 |
|  | 3 |  |  |  | -2 ± 1 | -5 ± 2 |

**Figure 7.**
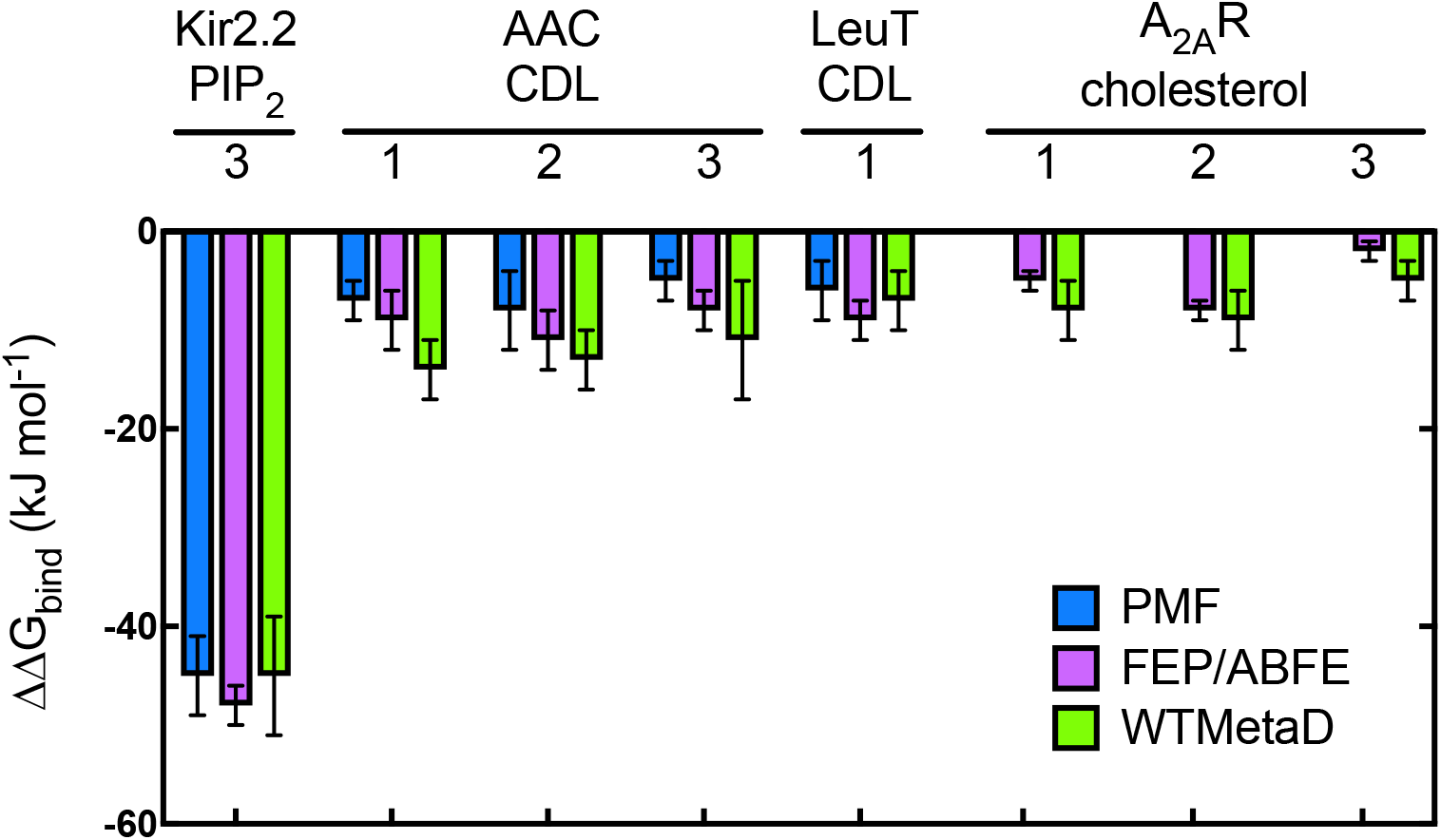
Comparison of different techniques. Bar chart showing ΔΔG_bind_ for the different protein-lipid systems described above. Shown are energies calculated with PMF (blue), FEP/ABFE (maroon) and WTMetaD (green). The error bars shown here are from the 200 rounds of bootstrap analysis for the PMF, standard deviations of 5 repeats for the FEP, or standard deviations of the energies for each site for the WTMetaD (green).

In addition to accuracy, it is of interest to compare the computational efficiency of the three methods, especially if they are to be employed in an automated pipeline (see e.g. ^31^) to characterise and compare protein-lipid interactions across a wide range of membrane protein structures. As can be seen from Table 2, the least computationally demanding technique is FEP/ABFE. This is because each alchemical pathway can be adequately described in 21 (PIP_2_), 29 (cholesterol) or 31 (CDL) windows, and 250 ns of simulation time per window is sufficient for good convergence (see Supporting Figures 2, 6 and 9). Running 5 independent repeats (resulting in ~25-40 μs simulation time in total) permits statistical analysis of the data. In fact, the data suggest that this cost could be further reduced to 150 ns per window with only 3 repeats (~9.5 μs in total; achievable in about 1 day with a mid to high range GPU), as this agrees well with the more extensively sampled data (e.g. −48 ± 2 kJ mol^-1^ vs −51 ± 1 kJ mol^-1^; Supporting Figure 13A). Note that these estimates do not account for the initial set of simulations that is required for the perturbation of each lipid species in bulk membrane, as this benchmark may be used for subsequent calculations.

**Table 2.** Computational cost of each technique as applied here. Reporting the simulation time used in each of the analysis measures here. Note that, as described in the Discussion, these values may overestimate the time required for convergence.

| technique | system applied to here | required simulation time |
| --- | --- | --- |
| PMF | Kir2.2, AAC, LeuT | 50-75 $\mu\text{s}$ |
| FEP | Kir2.2, AAC, LeuT | 25-40 $\mu\text{s}$ |
| ABFE | A<sub>2</sub>AR | 35 $\mu\text{s}$ |
| WTMetaD | Kir2.2, AAC, LeuT, A<sub>2</sub>AR | 120-190 $\mu\text{s}$ |

PMFs are generally less cost-efficient than FEP, taking at least 50 μs to converge. This is largely because equilibration of a lipid as it diffuses within a bilayer is relatively slow, meaning that each window needs to be simulated for 1 μs (see Supporting Figures 1, 5 and 8). In addition, a 0.05 nm window spacing from the bound state to bulk membrane usually requires 60 or more windows to get from the bound state to bulk. It should be noted, however, that a less frequent spatial sampling of 0.1 nm intervals, whilst resulting in a lower statistical certainty across the PMF, produces a similar estimation of ΔΔG_bind_ (for instance −45±2 kJ mol^-1^ vs −48±2 kJ mol^-1^; Supporting Figure 13B). In contrast to FEPs, however, PMFs offer a reaction coordinate in 1D space, which may be useful in certain cases. An example of this is for Kir2.2, where the PMF was able to detect the second, previously identified ^53^, binding site (Figure 3B).

Generally, WTMetaD is the most expensive technique, taking >100 μs to converge (see Supporting Figures 4, 7, 10 and 12), which is compounded by the requirement to read/write the Gaussian depositions to file, together taking ~3x longer to sample the same timescales as a standard MD simulation. However, in instances where a binding site is known, the simulations could be scaled back to focus upon the single binding site, saving considerable resources in the process. In addition, WTMetaD allows the mapping of the full 2D energy landscape around the target protein. This proved to be especially powerful for the A_2A_R and LeuT systems, where multiple asymmetric binding sites were revealed. For the Kir2.2-PIP_2_ system, the 2D landscapes were additionally compared to the PMF data through extraction of 1D coordinates from the data (see the Supporting Methods for details, and Supporting Figure 4F-G for data).

At present, we have compared these methods for estimation of lipid interaction free energies using the Martini ^22,23^ force field. We are aware that the free energies estimated are therefore an approximation due to smoothing of the free energy landscape, an inevitable consequence of coarse-graining. In the future, comparison of these data with corresponding atomistic simulations will be necessary, although achieving convergence of equivalent atomistic simulations is currently extremely challenging for systems of the size and complexity described here.

Finally, comparison of these methods to experimental analyses will be of particular importance. Currently, there have been only a few examples of experimentally-determined protein-lipid binding affinities: these include using nMS ^15^, SPR ^16^ and FRET ^29^. The majority of these approaches, however, involve both the protein and lipid being solubilised beforehand in detergent micelles. This will likely provide an inaccurate picture of the true energetics of lipid binding/unbinding in a lipid membrane, as the acyl tails will likely contribute substantially to the binding energies. Our work therefore highlights a pressing need for accurate measurement of lipid binding affinities to be applied to a number of well characterised membrane protein systems.

In summary, we have shown that three distinct methods for coarse-grained free energy calculations are able to provide robust estimates of the strength and specificity of lipid binding sites on membrane proteins. We envisage that these simulations can be readily performed, taking protein-lipid binding sites identified from structures, long equilibrium simulations (‘fingerprinting’) ^99,69^ or WTMetaD, and providing energies for specific lipid association in a semi-automated manner. The characterisation of lipid binding sites on integral membrane proteins offers the prospect of discovery of potential druggable allosteric sites on a wide range of membrane proteins, including those within the current membrane protein structural proteome of *ca*. 4,000 structures ^32^.

## Acknowledgements

RAC, PJS and MSPS are funded by the Wellcome Trust [208361/Z/17/Z] and the BBSRC [BB/P01948X/1, BB/I019855/1, BB/R00126X/1]. ONV and PJS are funded by the BBSRC [BB/P01948X/1]. Simulations were carried out in part on the ARCHER UK National Supercomputing Service (http://www.archer.ac.uk), provided by HECBioSim, the UK High End Computing Consortium for Biomolecular Simulation (hecbiosim.ac.uk), which is supported by the EPSRC [EP/L000253/1].

