## Supplement for "Insights into Membrane Protein-Lipid Interactions from Free Energy Calculations"

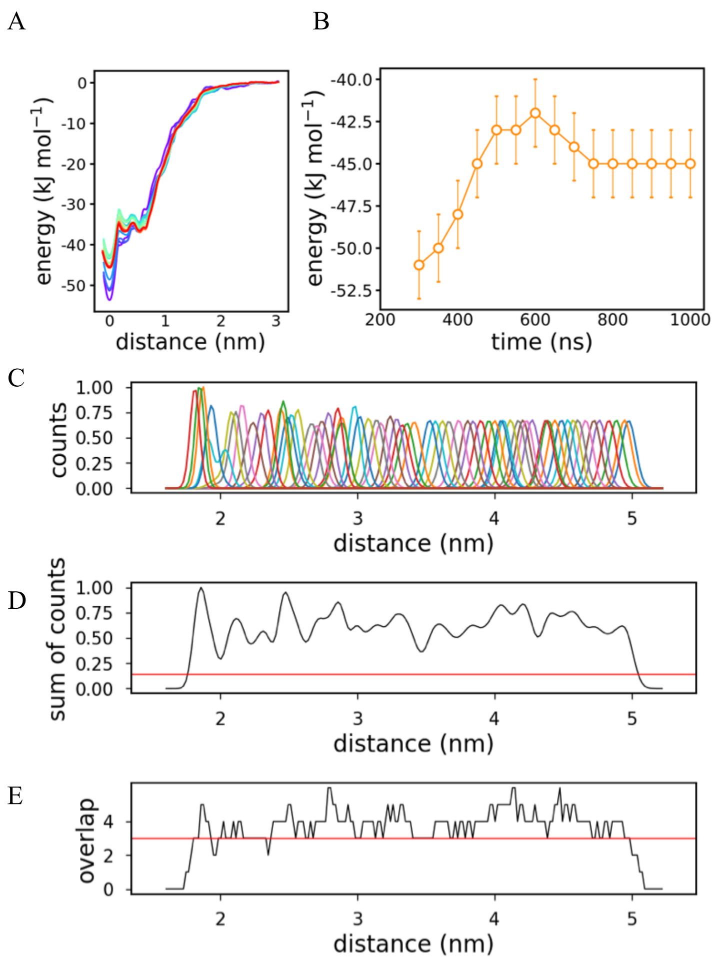


**Supporting Figure 1. Kir2.2 PMF convergence analysis**

A) Convergence analysis for the Kir2.2-$PIP_2$ PMF calculations. 200 ns were removed from the start of each window for equilibration purposes, before PMF profiles were built using increasing amounts of the simulation time, up to 1000 ns in total. The profiles were plotted for each run, coloured from purple (300 ns total) to red (1000 ns in total). Note that the red line is the same data as used for the main analyses.

B) Energy minima extracted from each of the traces in panel A, plotted as a product of simulation time analysed.

C) The raw histogram data produced from WHAM analysis. Each coloured histogram represents an independent umbrella sampling simulation.

D) Sum of the total counts along the reaction coordinate between Kir2.2 and $PIP_2$ as extracted from the histogram data in panel B. The data are normalised to 1. A line in red marks the threshold for sufficient sampling (0.2 of total counts).

E) The count of overlap histograms at each point along the reaction coordinate from the data in panel C, where only histograms which overlap at >10% of the total histogram height are counted. A red line marks the optimal overlap threshold of 3.

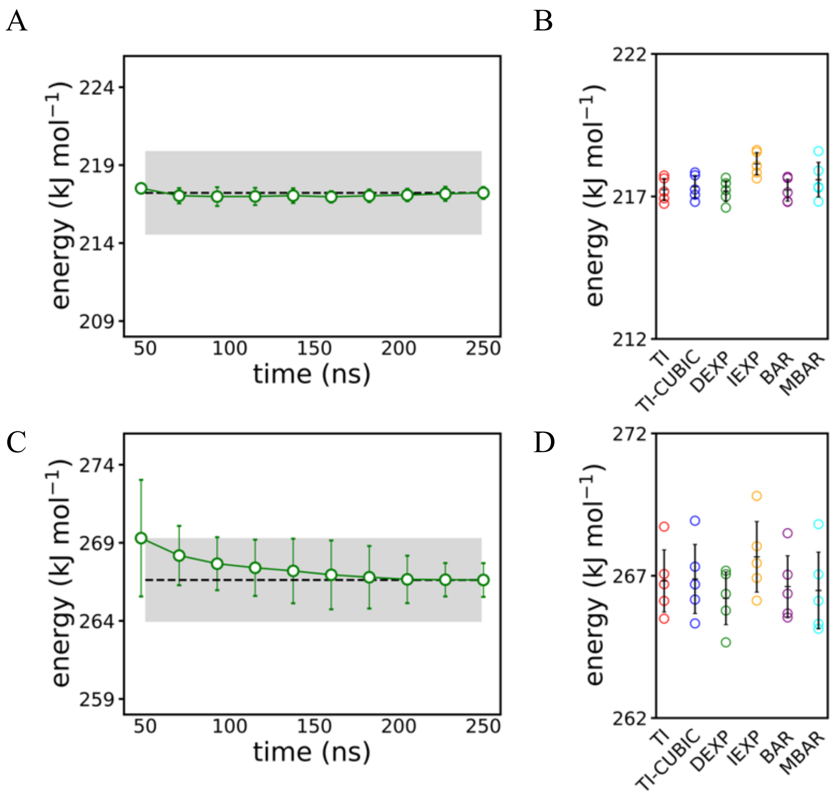


**Supporting Figure 2. Kir2.2 FEP convergence analysis**

A) Energies for perturbing a $PIP_2$ to POPC molecule free in a POPC membrane, as computed using MBAR over increasing amounts of simulation time. Here, the first 25 ns of each simulation has been discarded as equilibration. Plotted are the mean and standard deviation of 5 independent repeats, with a dotted line through the final mean value (250 ns). A grey bar marks the *RT* above and below the mean.

B) The computed energies for perturbing a $PIP_2$ to POPC molecule free in a POPC membrane following 250 ns of simulation, as calculated using different methods. TI=thermodynamic integration, TI-CUBIC=thermodynamic integration with a cubic spline, DEXP=deletion exponential averaging, IEXP=insertion exponential averaging, BAR=Bennett's acceptance ratio and MBAR=multistate Bennett's acceptance ratio.

C) As panel A, but showing data for perturbing a $PIP_2$ to POPC molecule whilst bound to the Kir2.2 channel.

D) As panel B, but showing data for perturbing a $PIP_2$ to POPC molecule whilst bound to the Kir2.2 channel.

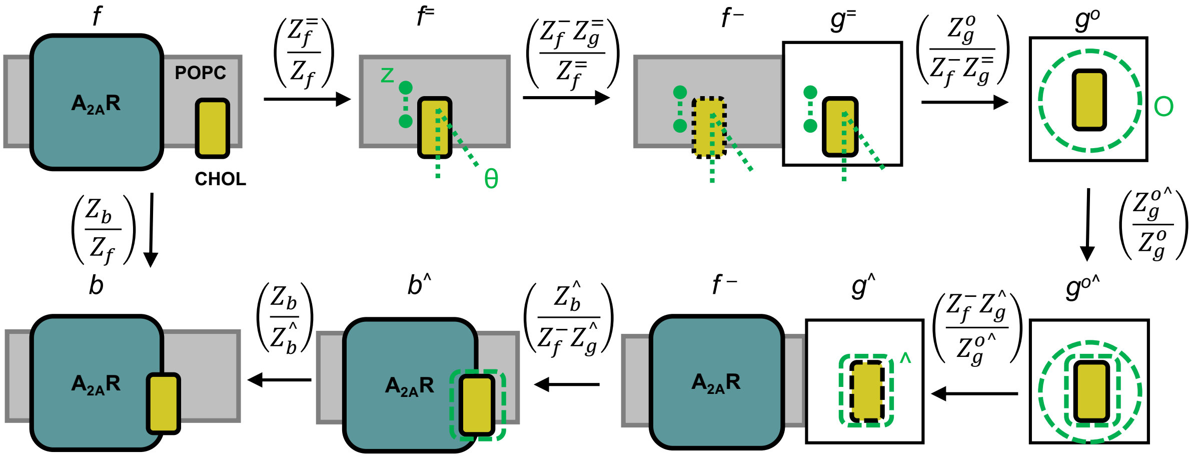


**Supporting Figure 3. Thermodynamic cycle for ABFE**

Shown is the thermodynamic cycle used for ABFE calculations of a target lipid to a generic lipid. The protein receptor is shown as a cyan square (labelled $A_{2A}R$), the bulk membrane as a gray square (labelled POPC, for its primary lipid constituent), and the target lipid shown in mustard (labelled chol), with a solid outline for its native state, and a dashed outline for its decoupled state. Restraints are shown in dotted green lines and labelled with the restraint name, with details of these outlined in the Supplementary Methods.

The top horizontal vertex represents the decoupling of the cholesterol from the membrane-only system. The right-hand vertex represents transferring the restraints method in the gas phase. The bottom horizontal vertex represents decoupling the cholesterol from the bound state. Above each arrow is shown the ratio of configuration partition functions (Z) for the step. For each configuration partition function, the restraint scheme imposed is shown as the upper indices (=, -, o or ^) as well as the lipid state on the lower indices (free='f', bound='b', gas phase='g'). These ratios are used to construct the final free energy calculation shown in Supplementary Equation 1 and 2.

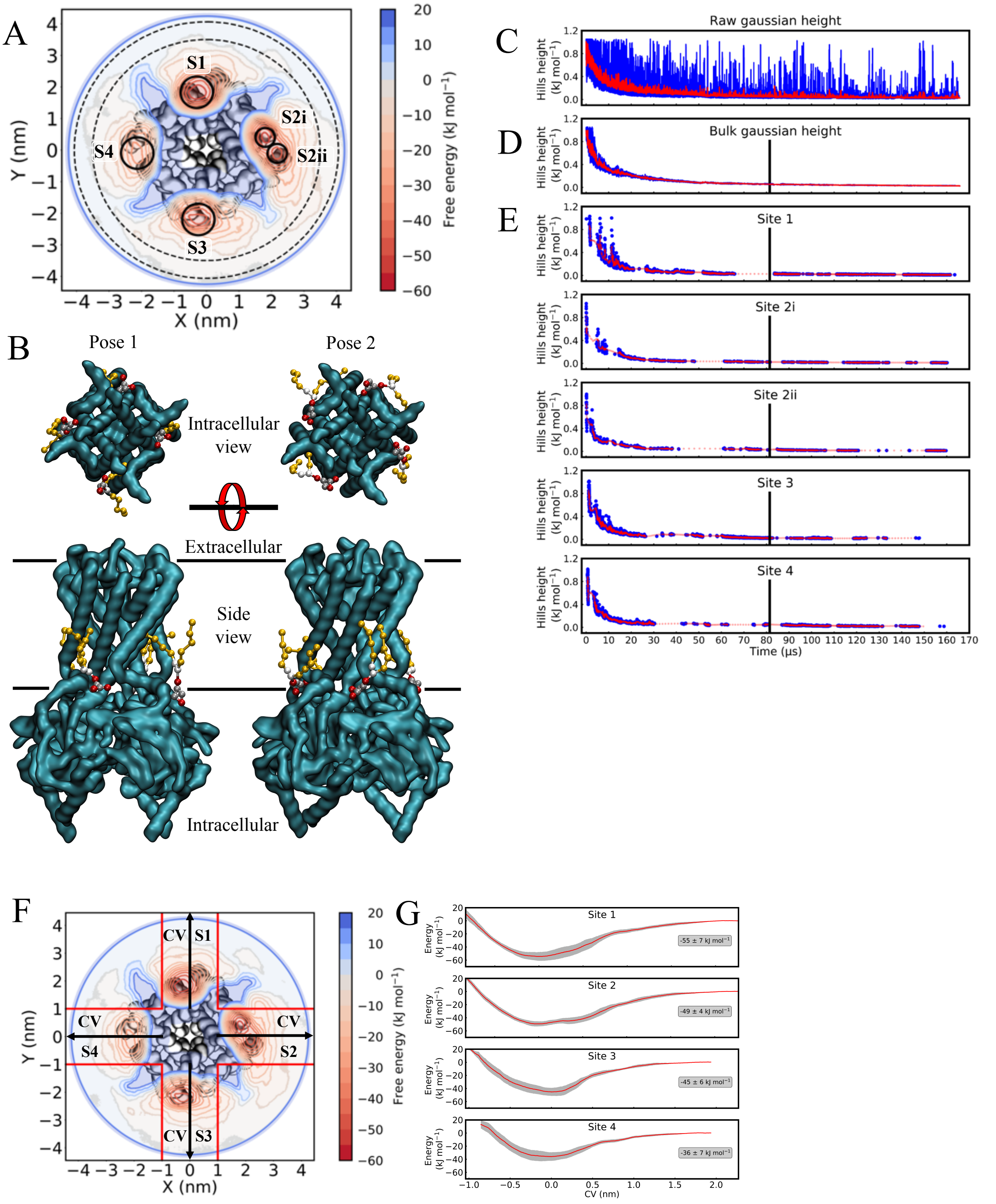


**Supporting Figure 4. Kir2.2 WTMetaD supporting data**

A) Shows the averaged 2D free energy landscape of Kir2.2 and $PIP_2$. The average free energy of the area between the dashed black lines was used as bulk. The area used to calculate each site is highlighted by the numbered circles on the energy landscape.

B) The four PIP2 binding sites highlighted by the WTMetaD simulations are shown from a Intracellular view and a side view. At each binding site two binding poses were recovered from the WTMetaD simulations referred to as pose one and two. Due to the symmetry of Kir2.2 only a single binding site has been shown. At sites 1, 3 and 4 it is not possible to differentiate between pose one and two in the

2D free energy landscape. The binding free energy is highlighted by each binding site and is reported in kJ $mol^{-1}$. The lipids are colour coded as acyl: yellow, aromatic: grey, glycerol: white and phosphate: red.
C) The raw Gaussian height of the WTMetaD run, only every 1 ns shown in blue with a running average of 10 shown in red.
D) The Gaussian height when the lipid is in the bulk environment. A black line denotes the time at which the bulk Gaussian is less than 5% of the maximum height which was discarded as equilibration.
E) The Gaussian height within 0.5 nm of each site is reported in blue every 1 ns with a running average of 10 shown in red.
F) Shows the averaged 2D free energy landscape of $PIP_2$ and Kir2.2. The energy minima every was recorded in 0.005 nm segments along the CV (black arrow) delimited by the red lines 1nm perpendicular to the CV.
G) Shown here is the average 1D energy landscape along the described CV. The energy landscape was taken every 250 ns after equilibration. Each data point was subsequently bootstrapped with resampling 10,000 times. The red line denotes the bootstrapped mean and the grey shading denotes the bootstrapped standard deviation. The energy minima for each landscape was referenced to 0 nm and the bulk was referenced to 0 kJ $mol^{-1}$. The reported energies for each site are taken from the energy minima (0 nm) whilst the errors are the sum of the standard deviation from the bulk and energy minima.

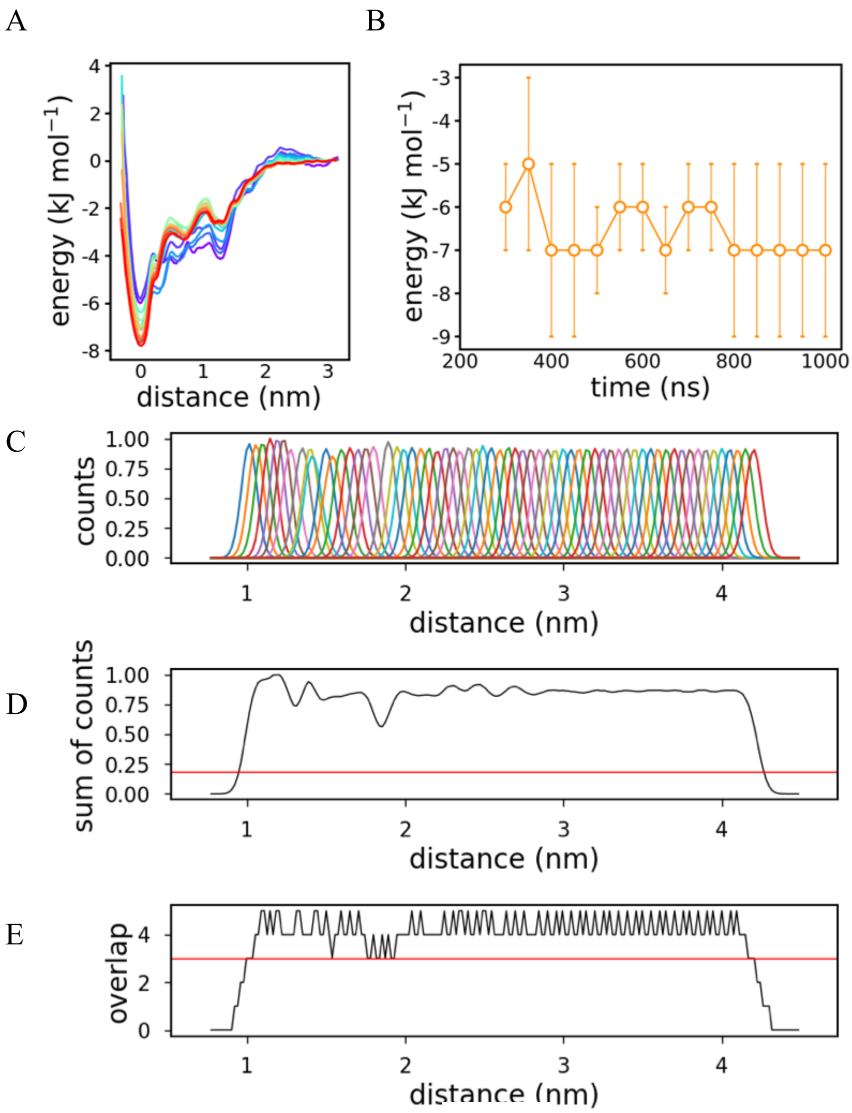


**Supporting Figure 5. AAC site 1 convergence analysis**

A) Convergence analysis for the AAC-CDL PMF calculations, as per Supporting Figure 1A

B) Energy minima extracted from each of the traces in panel A, plotted as a product of simulation time analysed.

C) The raw histogram data produced from WHAM analysis, as per Supporting Figure 1C.

D) Sum of the total counts along the reaction coordinate between AAC and CDL, as per Supporting Figure 1D

E) The count of overlap histograms at each point along the reaction coordinate, as per Supporting Figure 1E.

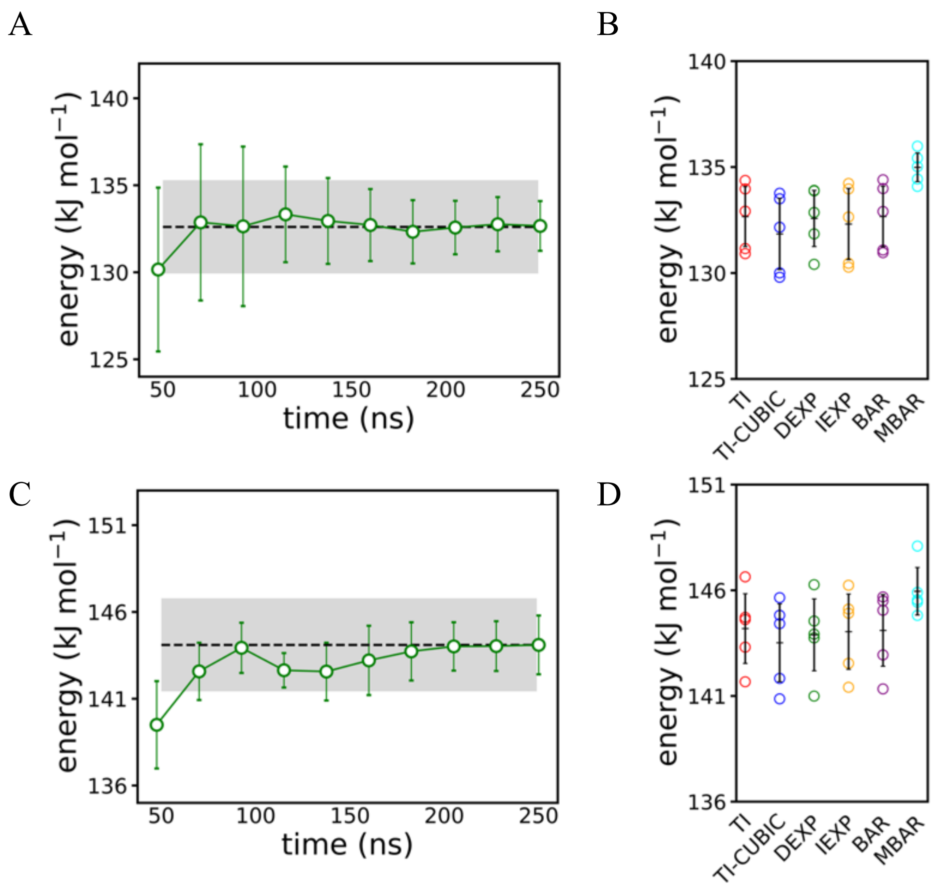


**Supporting Figure 6. AAC FEP convergence analysis**

A) Energies for perturbing a CDL to POPC molecule free in a POPC membrane, as computed using MBAR over increasing amounts of simulation time. Graph as per Supplementary Figure 2A.

B) The computed energies for perturbing a CDL to POPC molecule free in a POPC membrane following 250 ns of simulation, as calculated using different methods.

C) As panel A, but showing data for perturbing a CDL to POPC molecule whilst bound to AAC.

D) As panel B, but showing data for perturbing a CDL to POPC molecule whilst bound to AAC.

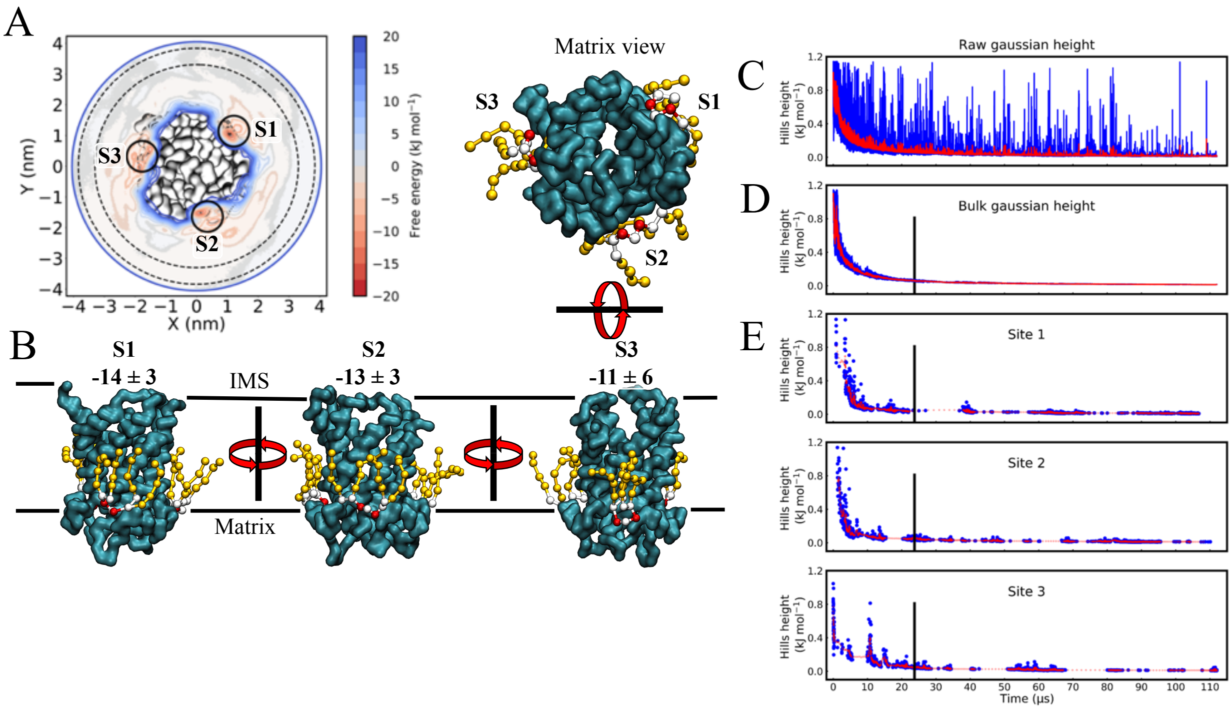


**Supporting Figure 7. AAC WTMetaD supporting data**

A) Shows the averaged 2D free energy landscape of AAC and cardiolipin. The average free energy of the area between the dashed black lines was used as bulk. The area used to calculate each site is highlighted by the numbered circles on the energy landscape.

B) The three binding sites highlighted by the WTMetaD simulations are shown as from a intracellular view and from either side. The binding free energy is highlighted by each binding site and is reported in kJ $mol^{-1}$.The lipids are colour coded as acyl: yellow, glycerol: white, phosphate: red.

C) The raw Gaussian height of the WTMetaD run, only every 1 ns shown in blue with a running average of 10 shown in red.

D) The Gaussian height when the lipid is in the bulk environment. Black line denotes the time at which the bulk Gaussian is less than 5% of the maximum height which was discarded as equilibration.

E) The Gaussian height within 0.5 nm of each site is reported in blue every 1 ns with a running average of 10 shown in red.

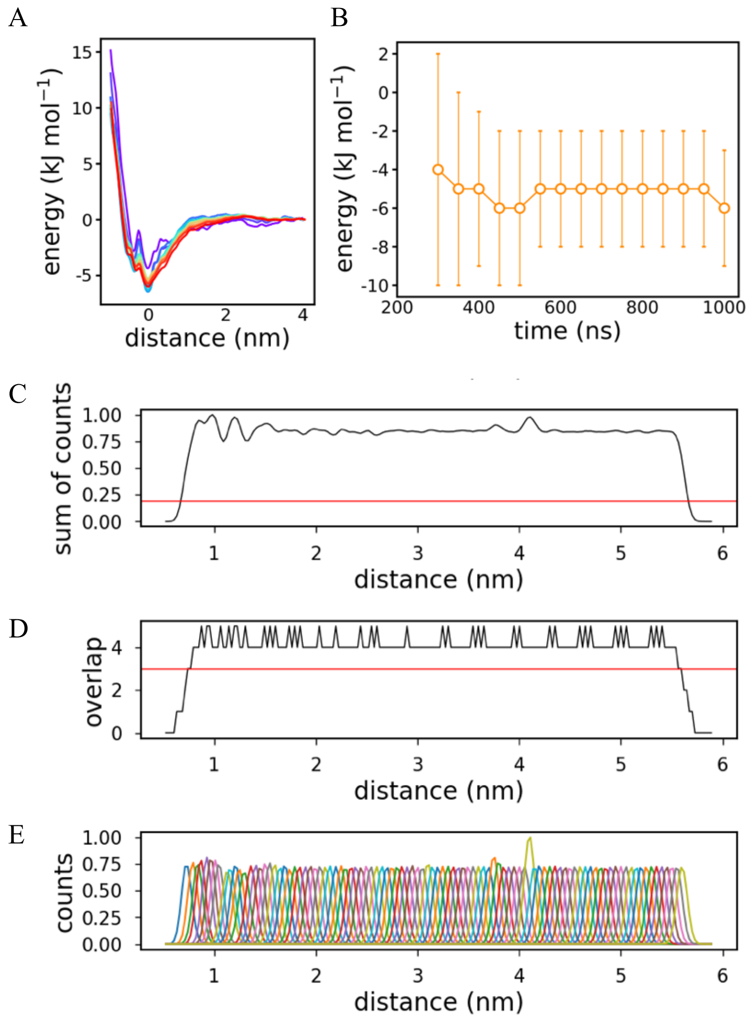


**Supporting Figure 8. LeuT PMF convergence analysis**

A) Convergence analysis for the LeuT-CDL PMF calculations, as per Supporting Figure 1A

B) Energy minima extracted from each of the traces in panel A, plotted as a product of simulation time analysed.

C) The raw histogram data produced from WHAM analysis, as per Supporting Figure 1C.

D) Sum of the total counts along the reaction coordinate between LeuT and CDL, as per Supporting Figure 1D

E) The count of overlap histograms at each point along the reaction coordinate, as per Supporting Figure 1E.

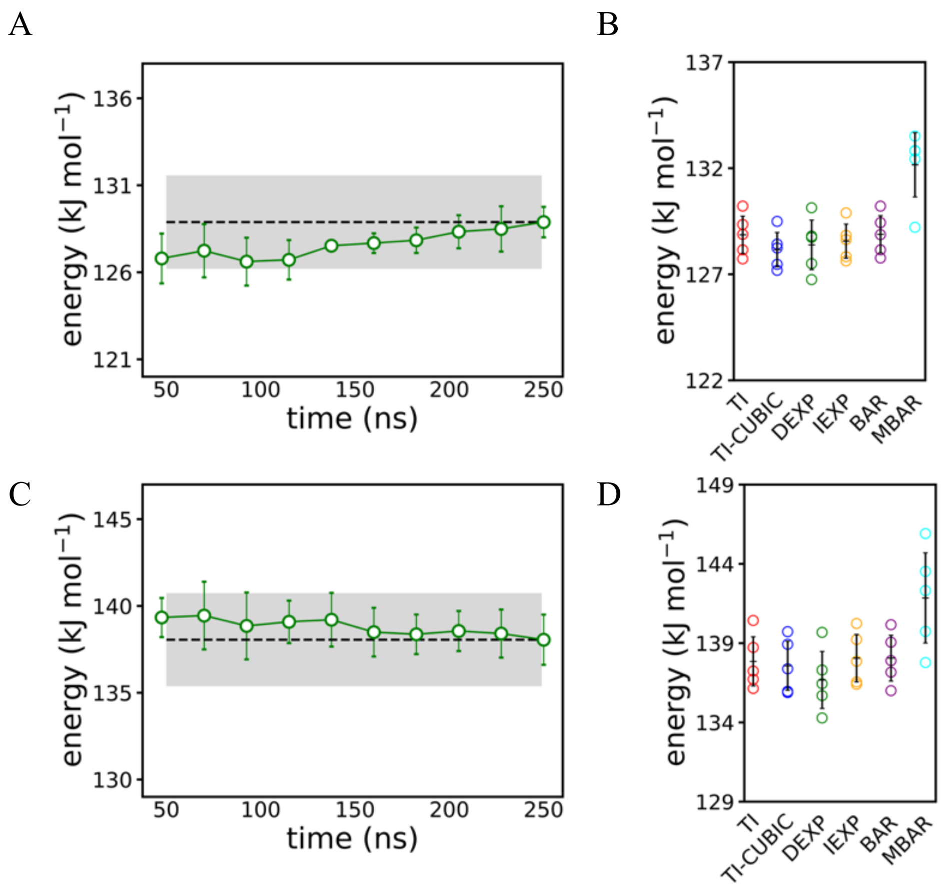


**Supporting Figure 9. LeuT FEP convergence analysis**

A) Energies for perturbing a CDL to POPE molecule free in a POPE membrane, as computed using MBAR over increasing amounts of simulation time. Graph as per Supplementary Figure 2A.

B) The computed energies for perturbing a CDL to POPE molecule free in a POPC membrane following 250 ns of simulation, as calculated using different methods.

C) As panel A, but showing data for perturbing a CDL to POPE molecule whilst bound to LeuT.

D) As panel B, but showing data for perturbing a CDL to POPE molecule whilst bound to LeuT.

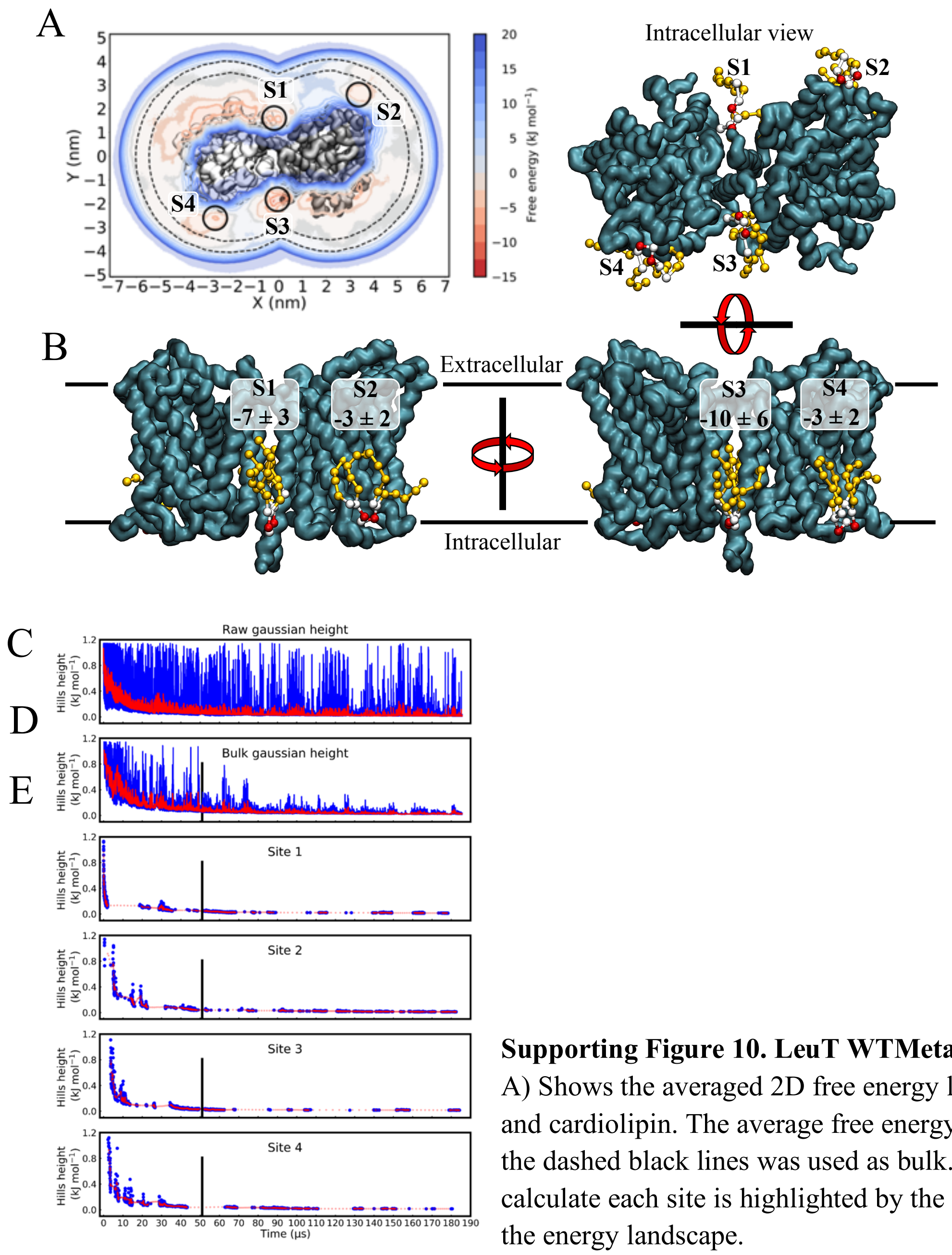


**Supporting Figure 10. LeuT WTMetaD supporting data**

A) Shows the averaged 2D free energy landscape of LeuT and cardiolipin. The average free energy of the area between the dashed black lines was used as bulk. The area used to calculate each site is highlighted by the numbered circles on the energy landscape.

B) The four binding sites highlighted by the WTMetaD simulations are shown as from a intracellular view and from either side. The binding free energy is highlighted by each binding site and is reported in kJ mol-1. The lipids are colour coded as acyl: yellow, glycerol: white, phosphate: red.

C) The raw Gaussian height of the WTMetaD run, only every 1 ns shown in blue with a running average of 10 shown in red.

D) The Gaussian height when the lipid is in the bulk environment. Black line denotes the time at which the bulk Gaussian is less than 5% of the maximum height which was discarded as equilibration.

E) The Gaussian height within 0.5 nm of each site is reported in blue every 1 ns with a running average of 10 shown in red.

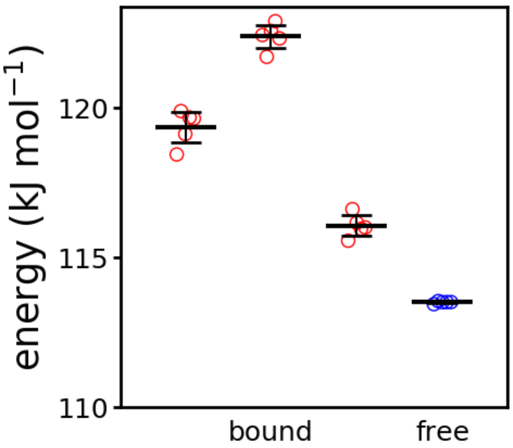


**Supporting Figure 11. $A_{2A}R$ ABFE supporting data**

FEP values for perturbation of cholesterol in each of the three $A_{2A}R$ sites (red) and free in the membrane (blue). Note that $\Delta\Delta G_{bind}$ cannot be directly computed from these values, as other components of the thermodynamic cycle in Supporting Figure 3 need to be accounted for.

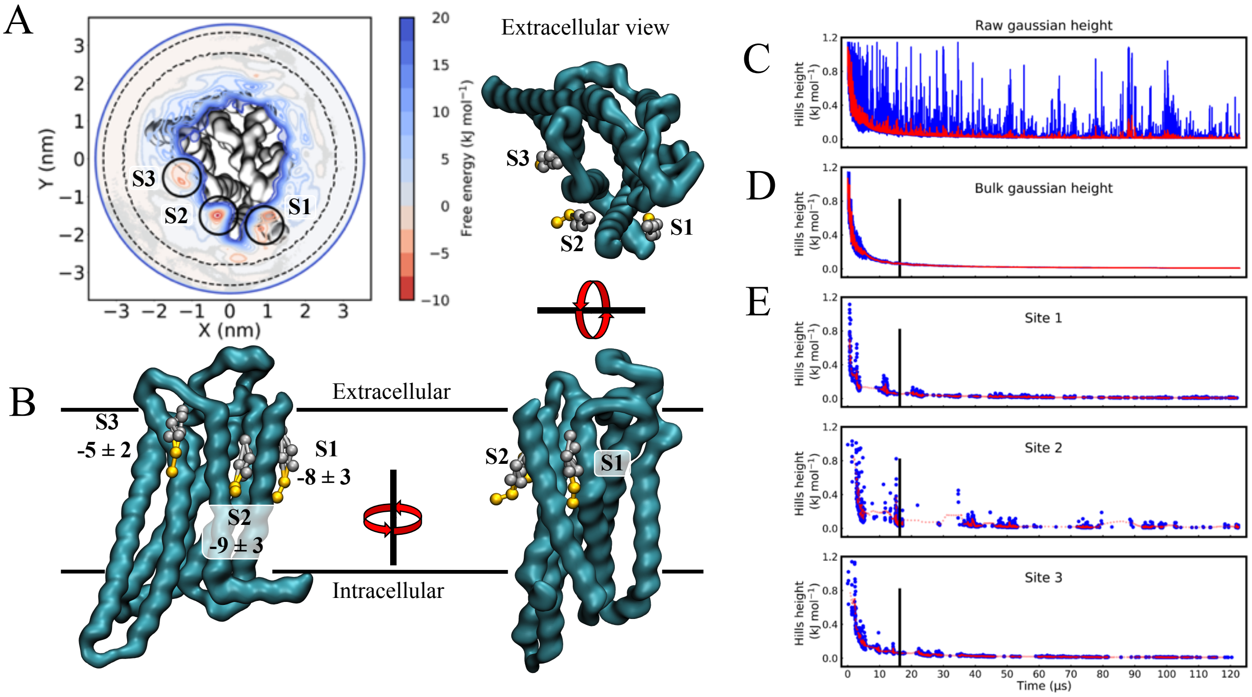


**Supporting Figure 12. $A_{2A}R$ WTMetaD supporting data**

A) Shows the averaged 2D free energy landscape of $A_{2a}AR$ and cholesterol. The average free energy of the area between the dashed black lines was used as bulk. The area used to calculate each site is highlighted by the numbered circles on the energy landscape.

B) The three cholesterol binding sites highlighted by the WTMetaD simulations are shown as from a extracellular view and from either side. The binding free energy is highlighted by each binding site and is reported in kJ $mol^{-1}$. The lipids are colour coded as acyl: yellow, aromatic: grey.

C) The raw Gaussian height of the WTMetaD run, only every 1 ns shown in blue with a running average of 10 shown in red.

D) The Gaussian height when the lipid is in the bulk environment. Black line denotes the time at which the bulk Gaussian is less than 5% of the maximum height which was discarded as equilibration.

E) The Gaussian height within 0.5 nm of each site is reported in blue every 1 ns with a running average of 10 shown in red.

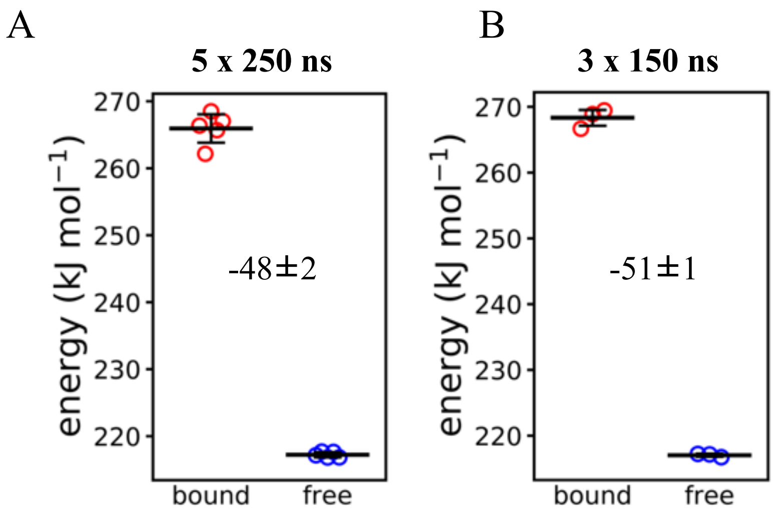


**Supporting Figure 13. Sampling frequency for FEP and PMF**

A) Comparison of FEP energies for perturbing a $PIP_2$ molecule to POPC either bound to Kir2.2 or free in the membrane. On the left is the analysis using the full 250 ns of each λ window, and with 5 repeats, representing ca. 26 µs and ca. 10 µs of simulation data.

B) Comparison of PMF profiles for $PIP_2$-Kir2.2 using either 0.05 or 0.1 nm spacing along the reaction coordinate.

$$\frac{Z_b}{Z_f} = \left(\frac{Z_f^=}{Z_f}\right)\left(\frac{Z_f^- Z_g^=}{Z_f^=}\right)\left(\frac{Z_g^o}{Z_f^- Z_g^=}\right)\left(\frac{Z_g^{o\wedge}}{Z_g^o}\right)\left(\frac{Z_f^- Z_g^\wedge}{Z_g^{o\wedge}}\right)\left(\frac{Z_b^\wedge}{Z_f^- Z_g^\wedge}\right)\left(\frac{Z_b}{Z_b^\wedge}\right)$$

**Supporting Equation 1. Calculation of binding energies from the thermodynamic cycle in Supporting Figure 3.**

The configuration partition function for each state is shown (Z), with the restraint scheme imposed shown (=, -, o or ^) as well as the lipid state (free='f', bound='b', gas phase='g').

$$\frac{Z_b}{Z_f} = \left(\frac{Z_f^=}{Z_f}\right)\left(\frac{Z_f^- Z_g^=}{Z_f^=}\right)\left(\frac{Z_g^o}{Z_g^=}\right)\left(\frac{Z_g^{o\wedge}}{Z_g^o}\right)\left(\frac{Z_b^\wedge}{Z_f^- Z_g^\wedge}\right)$$

**Supporting Equation 2. Modified form of Supporting Equation 1.**

Terms which cancel out or can be disregarded have been removed to create a modified form of the free energy function (as per Salari et al[1]). Terms 1 and 3 can be calculated analytically (see reference [1]). Term 2 is the FEP of the lipid in bulk membrane, term 4 is the restraint FEP in the gas phase, and term 5 is the FEP of the lipid when bound to the receptor.

$$\frac{Z_g^o}{Z_g^=} = \frac{2\pi r_R^3}{3z_R A^=}\frac{1}{1-\cos\theta_R}$$

**Supporting Equation 3. Calculation of second analytical term in Supporting Equation 2.**

How the second analytical term (term 3 in Supporting Equation 2) is calculated. Here, $r_R$ and $\theta_R$ are the values used for the distance and angle terms in the bulk lipid restraints, and $z_R$ is the distance used in the COM distance restraints in the gas phase.


(1) Salari, R.; Joseph, T.; Lohia, R.; Hénin, J.; Brannigan, G. A Streamlined, General Approach for Computing Ligand Binding Free Energies and Its Application to GPCR-Bound Cholesterol. *J. Chem. Theory Comput.* **2018**, *14* (12), 6560–6573. https://doi.org/10.1021/acs.jctc.8b00447.
